# GRK2-mediated AKT activation controls cell cycle progression and G2-checkpoint in a p53-dependent manner

**DOI:** 10.1101/2024.01.26.577358

**Authors:** Verónica Rivas, Teresa González-Muñoz, Ángela Albitre, Vanesa Lafarga, Federico Mayor, Petronila Penela

## Abstract

Cell cycle checkpoints, activated by stressful events, halt the cell cycle progression, and prevent the transmission of damaged DNA. These checkpoints prompt cell repair but also trigger cell death if damage persists. Decision-making between these responses is multifactorial and context-dependent, with the tumor suppressor p53 playing a central role. In many tumor cells, p53 alterations lead to G1/S checkpoint loss, rendering cell viability dependent on the strength of the G2 checkpoint through mechanisms not fully characterized. Cells with a strong pro-survival drive can evade cell death despite substantial DNA lesions. Deciphering the integration of survival pathways with p53-dependent and -independent mechanisms governing the G2/M transition is crucial for understanding G2 arrest functionality and predicting tumor cell response to chemotherapy. The serine/threonine kinase GRK2 emerges as a signaling node in cell cycle modulation. In cycling cells, but not in G2 checkpoint-arrested cells, GRK2 protein levels decline during G2/M transition through a process triggered by CDK2-dependent phosphorylation of GRK2 at the S670 residue and Mdm2 ubiquitination. We report now that this downmodulation in G2 prevents the unscheduled activation of the PI3K/AKT pathway, allowing cells to progress into mitosis. Conversely, higher GRK2 levels lead to tyrosine phosphorylation by the kinase c-Abl, promoting the direct association of GRK2 with the p85 regulatory subunit of PI3K and AKT activation in a GRK2 catalytic-independent manner. Hyperactivation of AKT is conditioned by p53’s scaffolding function, triggering FOXO3a phosphorylation, impaired cyclin B1 accumulation, and CDK1 activation, causing a G2/M transition delay. Upon G2 checkpoint activation, GRK2 potentiates early arrest independently of p53 through AKT activation. However, its ability to overcome the G2 checkpoint in viable conditions depends on p53. Our results suggest that integrating the GRK2/PI3K/AKT axis with non-canonical functions of p53 might confer a survival advantage to tumor cells.

## INTRODUCTION

Cell cycle progression is tightly regulated by various mechanisms or checkpoints to ensure the orderly and faithful sequence of replication and division events (1, 2). These same regulatory processes are often exploited as targets for conventional anti-tumoral chemotherapies. The cell cycle checkpoints temporarily halt proliferation of the cells exposed to genotoxic compounds in response to the activation of the cellular DNA damage response (3, 4, 5). This pathway orchestrates diverse mechanisms for the repair and survival of cells (6). Instead, if the extent of damage surpasses the cells’ capacity for repair, it can result in cell death (7). Weakness in such repair mechanisms, particularly during the G1 or G2 phases, can lead to accumulation of genetically unstable cells over time, thereby contributing to cancer pathogenesis (1).

In many cancer cells, the G1/S checkpoint, which prevents the replication of damaged DNA, is compromised due to mutations in p53, a key regulator of the cell cycle and tumor suppressor (8, 9). As a result, their response to genotoxic agents primarily relies on the robustness of the G2 and S checkpoints (1). This dependence presents opportunities for novel treatments, as combining genotoxic agents with drugs designed to disrupt such cell cycle checkpoints might trigger synthetic lethality (10). In this context, the elimination of the G2 checkpoint sensitizes p53-deficent cells to chemotherapeutic-induced cell death (11). Cells passing through the G1 phase with unrepaired DNA will undergo mitotic catastrophe and apoptosis (12). Nevertheless, cells equipped with strong pro-survival mechanisms can evade cell death, reassuming the cell cycle and contributing to the promotion of genomic instability.

The G2 checkpoint employs redundant pathways to restrain the activation of CDK2/cyclin B1 complex, promoting the inactivating action of the tyrosine kinase Wee1 and the suppression of the activating phosphatase CDC25C (13). While the G2 checkpoint can be engaged in the absence of p53, this factor contributes to both the initial and long-term maintenance of G2 arrest (14, 15, 16). Additionally, the AKT signalling pathway can influence G2 checkpoint activation in opposite directions in a cell type and context-dependent manner (17, 18, 19, 20). Understanding the effects of PI3K/AKT in the G2 checkpoint and cell survival under genotoxic stress, and the interplay between PI3K/AKT and p53-dependent mechanisms is crucial for determining whether a cell will undergo a transient or prolonged G2 arrest in response to DNA damage, and for evaluating the efficacy of current chemotherapeutic drugs.

In this context, the G protein-coupled receptor kinase GRK2 has been suggested as a crucial player in coordinating cell growth and the PI3K/AKT pathway in response to mitogenic extracellular signals (21)) and regulating cell cycle progression dynamics (22, 23, 24). While GRK2 is primarily recognized for its role in modulating numerous G protein-coupled receptors (GPCRs) (25, 26), it also modulates additional signalling pathways through phosphorylation or scaffolding interactions with various non-GPCR cellular partners (27). This multifaceted role of GRK2 is very relevant for the proper and timely progression of the cell cycle (27).

In normal cell cycle progression, GRK2 is downregulated at the G2/M transition through phosphorylation at Ser670 by CDK2-Cyclin (22). This phosphorylation event allows the subsequent binding of the prolyl-isomerase Pin1, leading to the conjugation of polyubiquitin chains by Mdm2 for kinase degradation (22, 24). This degradation is necessary to regulate centrosome separation mediated by the EGF/MST2/NEK2 pathway to ensure the formation of mitotic spindles of the appropriate length (24). Interestingly, when the G2 checkpoint is activated and the cell cycle arrested due to the presence of DNA damaging agents like doxorubicin, GRK2 protein levels are actively stabilized, which helps limit the activation of p53 and the apoptotic response in G2/M arrested cells (22). However, how the genotoxic-induced stabilization of GRK2 impacts target proteins related to G2 checkpoint activation and maintenance, as well as its dependence on p53 functionality, remains to be elucidated. Of note, the enforced stabilization of GRK2 induces per se a G2 cell cycle delay in HeLa cells, although the mechanism underlying this effect is unknown.

In this study, we examined how GRK2 levels and p53 expression influence the G2/M transition of normal cycling cells and the proficiency of the G2/M checkpoint. We uncover that G2 phase-stabilized GRK2 acts as a scaffolding protein for AKT activation via c-Abl1 phosphorylation and interaction with the p85 regulatory subunit of class IA PI3K. The GRK2/PI3K/AKT signalosome, along with p53 transactivation-independent activities, contributes to the robustness of G2 checkpoint and survival, with potential implications for the chemotherapeutic response of tumor cells.

## RESULTS

### Stabilization of GRK2 in cycling cells results in delayed G2 phase, triggering a parallel PI3K-dependent activation of AKT

We have previously demonstrated that the phosphorylation status of GRK2 affects its protein stability in the context of the cell cycle, thereby altering the kinetics of cell cycle progression. The phosphorylation of GRK2 at Ser670 by CDK2-Cyclin A and the subsequent binding of the prolyl-isomerase Pin1 are required for transient GRK2 proteasomal degradation at the G2/M transition in HeLa cells (22). Interestingly, the expression of a mutant unable to be phosphorylated at this residue (GRK2-S670A) completely prevents GRK2 down-modulation and delays cell cycle progression though the G2 phase (22). A similar protracted G2/M transition occurs in the presence of a catalytically inactive GRK2-K220R mutant, which displays an inherently retarded degradation (22).

In search of mechanisms underlying such G2 phase delay, we investigated signalling pathways known to be involved in the G_2_/M transition and displaying functional connections with GRK2. GRK2 serves as a scaffold for the Raf/MEK/ERK, cascade (28, 29), whose activation fluctuates during the cell cycle, decreasing as cells progress through the G2 phase (30). Untimely activation of MEK/ERK pathway during G2 may result in a marked delay in cell cycle progression (31), and impaired activation of ERK1/2 or MEK1 is known to interfere with mitotic progression (30). In HeLa cells expressing GRK2-S670A (HeLa-A1) or GRK2-K220R (HeLa-K1) proteins, activation of ERK1/2 (Suppl.Fig.S1A) and MEK1 (Suppl.Fig.S1B) was similar to parental HeLa cells, showing a decrease in G2 followed by an increase between 10 and 14h post-release from a G1/S arrest. Despite no alterations in ERK or MEK1 activity were detected, progression into mitosis was clearly delayed in HeLa-A1 and HeLa-K1 cells. This delay was evidenced by the defective accumulation of Cyclin B1 and impaired tyrosine dephosphorylation of CDK1, both of which are required for the activation of the Cdk1/cyclin B complex (Suppl.Fig.S1C).

On the contrary, PI3K/AKT stimulation pattern was markedly affected in HeLa-A1 and HeLa-K1 cells. Whereas in parental cells, in line with previous reports a modest peak of AKT phosphorylation was detected within 2h-4h of release from a G1/S arrest, with minimal activity in the late G2 phase (Fig.1A). This pattern was maintained upon expression of extra wt-GRK2 levels (Suppl.Fig.S2A), a condition that did not alter normal cell cycle progression, as overexpressed kinase downmodulates in G2 similar to the endogenous (22). In contrast, the presence of degradation-defective GRK2-S670A or GRK2-K220R proteins led to a robust and maintained increase in PI3K-dependent phosphorylation of AKT, as demonstrated by the complete prevention of its activation with the PI3K inhibitor LY294002 (Fig. 1A-B). Of note, AKT activation in G2 was also higher in HeLa cells stably expressing a C-terminal construct of GRK2 (aa 436-689) (Suppl.Fig.S2B), that competes the interaction of CDK2/Pin1 with GRK2, resulting in the stabilization of the endogenous protein and a delay in G2 phase progression (Penela et al., 2010). Since similar results were observed in HeLa-A1, HeLa-K1 and HeLa-GRK2_436-689_ cells, we inferred that the common molecular factor triggering AKT activation could be either the accumulation of extra GRK2 protein or the impairment of GRK2 kinase activity reported for these mutants (27, 32). However, no increased phosphorylation of AKT during the G2 phase was detected in HeLa cells expressing a specific GRK2 shRNA construct expected to reduce catalytic activity as kinase protein levels are decreased by approximately 70% (Suppl.Fig.S2C). This suggest that a scaffolding function of GRK2 protein, regardless of its activity status, can trigger abnormal cell cycle activation of AKT.

**Figure 1.**
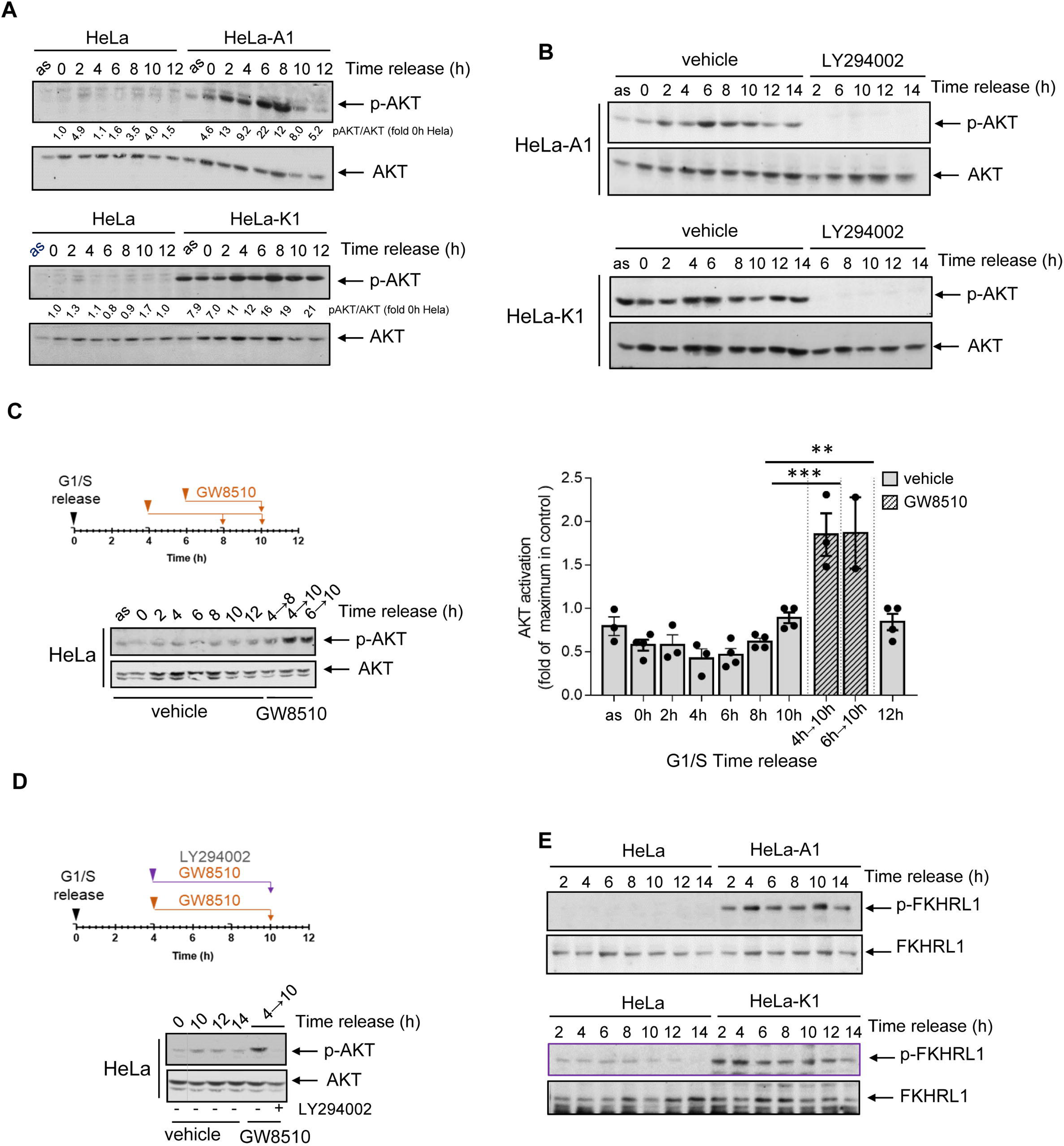
Impaired GRK2 downmodulation during G2 progression stimulates the PI3K/AKT signalling pathway. Expression of degradation-defective GRK2 proteins or interference with the cell cycle modulators of GRK2 degradation causes a strong PI3K-dependent activation of AKT during G2 progression. Control and HeLa cells with extra GRK2-S670A (HeLa-A1) or GRK2-K220R (HeLa-K1) were synchronized at the G1/S boundary and released into the cell cycle as detailed in Methods in the absence (A) or presence (B) of 10µM LY294002 added 4h post-released and maintained for the indicated times. Whole-cell lysates were analyzed by immunoblotting using either anti-AKT and anti-pS473-AKT antibodies. (C, D) Inhibition of CDK2 promotes the activation of AKT during G2 in a PI3K-dependent manner. HeLa cells synchronously entering the cell cycle were incubated with vehicle or 7µM GW8510 alone (C) or in the presence of LY294002 (D) for the indicated times. The phosphorylation status of AKT was assessed as above, and normalized data by total AKT were represented as fold-change with respect to maximal AKT activation in vehicle control cells. Data are mean ± SEM, n = 2–4 independent determinations. (E) G2-activated AKT in cells overexpressing degradation-defective GRK2 proteins is paralleled by an increase in the phosphorylation of the AKT substrate FKHRL1. Whole-cell lysates of HeLa-A1, HeLa-K1 and control HeLa cells progressing synchronously into G2 were analyzed by immunoblotting using anti-FHKRL1 and anti-pT32-FHKRL1 antibodies. Representative blots of at least 2-3 independent experiments are shown. Dashed lines indicate removal of lanes within the same blot

Consistently, treatment of parental HeLa cells with the CDK2 inhibitor GW8510, which prevents the phosphorylation of GRK2 by CDK and abolishes the decay of endogenous GRK2 (22), resulted in a significant increase in phosphorylated AKT protein 10h after release from a G1/S arrest (Fig.1C), with the involvement of PI3K activity (Fig.1 D).

Notably, a delayed entry into mitosis triggered by the impaired downregulation of PI3K activity during G2 has been linked to defective Forkhead-dependent transactivation of cyclin B (33). Consistent with this, significantly increased AKT-dependent phosphorylation of endogenous FKHRL1 (FOXO3a) was observed in HeLa-A1 and HeLa-K1 cells at all times following release from G1/S arrest (Fig.1E). This paralleled the impaired upregulation of Cyclin B1 and CDK1 activation (Suppl.Fig.S1C), while phospho-FKHRL1 was barely detected in control cells (Fig.1E), where protein Cyclin B1 accumulated timely (Suppl.Fig.S1C).

Taken together, these results strongly suggest that under normal conditions, the degradation of CDK2-phosphorylated GRK2 restrains the activation of the PI3K/AKT pathway in G2 and prevents the AKT-mediated nuclear exclusion of Forkhead transcription factors, which, in turn, allows for FOXO3a-dependent cyclin B synthesis increase (33).

### GRK2 directly associates with the p85 regulatory subunit of PI3K in an Abl-mediated tyrosine phosphorylation-dependent manner, leading to AKT activation and G2 delay

The above results suggested that increased scaffolding GRK2 activity linked to impaired downregulation of the protein promotes untimely PI3K activation during G2. GRK2 has been reported to interact with the catalytic subunits of the class I PI3K isoforms p110γ and p110β (34). Therefore, we investigated whether the GRK2-S670A mutant displayed a differential association with endogenous PI3K in cells progressing through the cell cycle. Since different p110 subunits form functional heterodimers with common p85 regulatory subunits, we used an antibody directed against the p85 subunit to detect the presence of GRK2-associated PI3K. We observed a clear GRK2/ p85-PI3K association in cells expressing GRK2-S670A (but not wt-GRK2), which correlated with AKT activation and the prevention of Cyclin B upregulation (Fig.2A).

**Figure 2.**
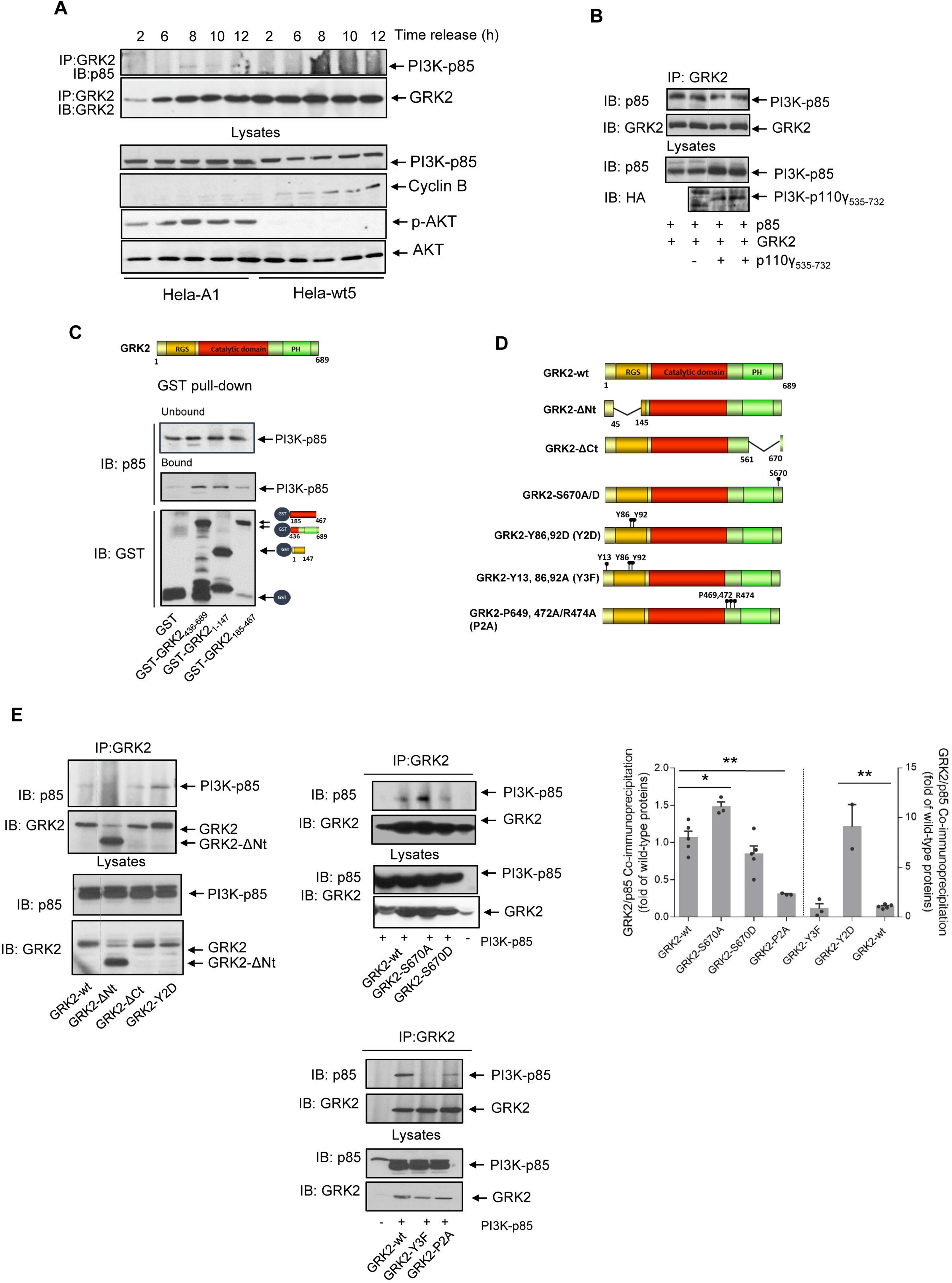
Abnormally accumulated GRK2 protein interacts with the regulatory p85 subunit of PI3K in G2 phase through N-terminal phosphorylated tyrosine residues and C-terminal proline-box. (A) Downmodulation-defective GRK2-S670A protein but not degradation-competent wt-GRK2 co-immunoprecipitated with p85 in cells progressing through G2 in correlation with improper ATK activation and impaired G2/M transition. HeLa cells stably expressing GRK2-wt (HeLa-wt5) or GRK2-S670A (HeLa-A1) were synchronized at the G1/S boundary and released into G2 for the times indicated. Cellular lysates were processed for GRK2 immunoprecipitation, and immune complexes were analyzed with an anti-p85 antibody as described in Methods. After stripping, the presence of total GRK2 was immunodetected in the same gels. Levels of p85 protein, cyclin B1, pAKT, and total AKT were analyzed in whole lysates. (B) The association of GRK2 with p85 was not mediated by PI3K-p110 subunit. Overexpression of the GRK2-binding PIK domain of PI3K-p110 does not compete for p85 binding to GRK2. HeLa cells were co-transfected with GRK2 and p85 in the presence or absence of an HA-tagged construct of the PIK domain (PI3K-p110γ_535-732_) and processed for GRK2 immunoprecipitation and detection of associated p85. The input level of the expression of p85 and PIK domain protein in cell lysates was assessed by immnunoblotting. (C) Recombinant p85 was incubated with GST alone or the indicated GST-GRK2 fusion proteins as described in Methods. Proteins bound to Glutathione-sepharose beads were immunodetected with specific anti-p85 and anti-GST antibodies. The GST pull-down blots are representative of 3 separate experiments. (D-E) Characterization of the GRK2 motifs involved in the interaction with p85. The co-immunoprecipitation of p85 was analyzed as above with several truncated and site-point GRK2 mutants depicted in D. HeLa cells were transiently co-transfected with p85, GRK2wt, deletion GRK2 mutants (GRK2-ΔNt, GRK2 1-147, GRK2ΔCt), GRK2 mutants with altered tyrosine (GRK2-Y2D, GRK2Y3F) or serine (GRK2-S670A, GRK2-S670D) phosphorylation, or the proline-box mutant GRK2-P2A (E). GRK2/p85 coimmunoprecipitation is based on the integrity of the kinase proline box and fostered by phosphorylation of tyrosine 86 and 92 within the RH domain of GRK2. Data (mean ± S.E.M of 3-5 independent experiments) indicate the amount of co-precipitated p85 protein, normalized by immunoprecipitated GRK2 and referred to the association displayed by wild-type proteins *, p< 0.05, ** p< 0.01 compared to GRK2wt/p85 association. Representative blots are shown. Dashed lines indicate removal of lanes within the same blot

Interestingly, while previous reports indicated a direct interaction between GRK2 and p110 catalytic subunits (35), our co-immunoprecipitation assays of co-expressed GRK2, p85 or p110α tagged proteins revealed a more consistent and readily association between GRK2 and p85. We observed that the co-immunoprecipitation of GRK2 with p85 remained unaffected in the presence of a construct containing the PIK domain of p110 (p110γ_535-732_) (Fig. 2B), which had initially been identified as the GRK2-interacting domain for class I PI3K (35). These results suggested that GRK2 could also bind to PI3K independently of p110, via its association with the p85 subunit.

The SH3 and SH2 domains of p85 allow the recruitment of various signalling proteins bearing Pro-rich motifs and phosphotyrosine residues, respectively, which are essential for regulating PI3K activity and membrane targeting. Interestingly, GRK2 possesses a proline-rich motif in the C-terminal region (^469^PLIPPR^474^) and several tyrosine residues in the RH domain (Y13,Y86,Y92), which can be phosphorylated by tyrosine kinase receptors (36) as well as by tyrosine kinases of the c-Src family (37). Pull-down assays using recombinant p85 and GST-GRK2 constructs encompassing different kinase domains revealed that p85 preferentially bound to GST-GRK2_1-147_ and GST-GRK2_436-689_, but not to GST alone or the GST-GRK2_185-467_ region which lacks the proline-rich motif (Fig.2C). These results suggested a multi-domain interaction between GRK2 and p85, involving the RH domain and the C-terminal portion of GRK2. Consistently, a GRK2-ΔNt mutant lacking most of the RH domain (GRK2_Δ50-145_) did not interact with p85 *in situ* (Fig.2D-E, left panel), while a protein construct bearing the entire RH domain (aa 1-147) did (Fig.2E). This interaction seems to be regulated by tyrosine phosphorylation, as it was notably enhanced in the presence of a GRK2 mutant that mimics permanent phosphorylation at tyrosines 86 and 92 (Fig.2E). In contrast, GRK2 mutated at the proline-box (GRK2-P2A) was unable to associate with p85, while a C-terminal deletion GRK2-ΔCt mutant, still bearing the proline-box (aa 467-479), was able to support p85 binding. Moreover, this proline-box can sustain a “basal” association with p85 in the absence of tyrosine phosphorylation since a GRK2 mutant defective in tyrosine phosphorylation displays the same binding as wt GRK2.

Overall, our results suggest that the RH domain and the C-terminal proline-box of GRK2 can mediate the binding to the SH2- and SH3-domains of p85, either independently or in combination.

We then examined whether the phosphorylation of GRK2 at Ser670 could influence p85 binding. The association of GRK2-S670D was not significantly different from that of the GRK2-wt protein, whereas the association of GRK2-S670A was increased (Fig.2E, right panel). Thus, the phosphorylation of S670 itself does not appear to be a key factor for “keeping away” p85-PI3K from GRK2. Instead, S670 phosphorylation is crucial for triggering Pin1 recruitment and GRK2 degradation (22, 24), ultimately reducing the pool of available GRK2 protein for binding to p85. This notion is consistent with the observation that the removal of aa 561-670 had no effect on the GRK2/p85 interaction, suggesting that neither this region nor its phosphorylation influences the conformational availability of the nearby proline-box motif. Hence, the increased association of the GRK2-S670A mutant with p85 during G2 progression must involve another mechanism.

Interestingly, we observed that the mutant GRK2-S670A protein, but not wt-GRK2, displays increased tyrosine phosphorylation as cells progress through G2/M (Fig.3A). This event parallels its enhanced association with p85 (Fig.2A). In an attempt to identify the tyrosine kinase responsible for this phosphorylation event, synchronized cells were treated with specific inhibitors for either c-Src or EGF receptor activities, since both kinases have been shown to phosphorylate GRK2 (36, 38). However, neither of these inhibitors affected the tyrosine-phosphorylation status of GRK2-S670A (Suppl.Fig.S3A). Interestingly, using the purified recombinant GRK2 protein as bait for SH3 domain binding, we identified several partner proteins in a peptide array that may potentially interact with the proline-rich sequence carried by GRK2 (Suppl.Fig.S3B). Besides c-Src, a well-known tyrosine kinase for GRK2 which detection validates the overlay assay (39), these partners included the Phosphoinositide-3-Kinase Regulatory Subunit 1 (p85α), consistent with our previous results (Fig. 2), as well as the non-receptor tyrosine kinase Abl2.

**Figure 3.**
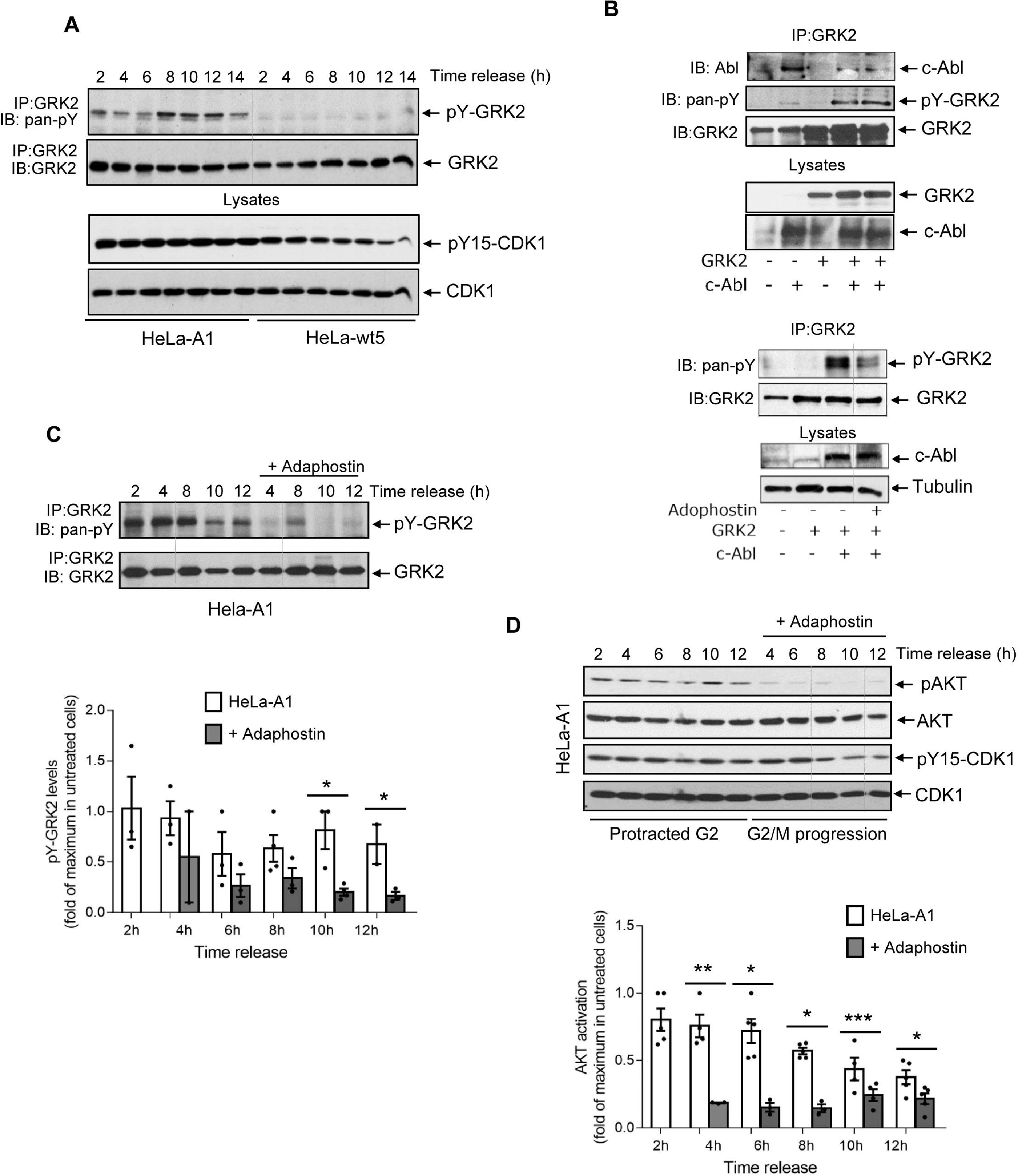
The c-Abl tyrosine phosphorylation of accumulated GRK2 in G2 is required for the activation of AKT. (A) The extent of tyrosine phosphorylation of GRK2-S670A increases during G2. HeLa cells stably expressing GRK2-wt (HeLa-wt5) or GRK2-S670A (HeLa-A1) were synchronized at the G1/S boundary for subsequent release, and immunoprecipitates of GRK2 analyzed with an anti-phosphotyrosine monoclonal antibody. After stripping, the presence of total GRK2 was immunodetected in the same gels. (B) c-Abl1 interacts with and increases the tyrosine phosphorylation of GRK2. HeLa cells were transiently transfected with GRK2 WT in the presence or absence of c-Abl1, and cellular lysates were subjected to immunoprecipitation with anti-GRK2, as detailed in Methods. Both GRK2 co-immunoprecipitated c-Abl1 protein and tyrosine phosphorylation of GRK2 (pY-GRK2) in the immunocomplex, in the absence or presence of 50 µM adaphostin, a catalytic inhibitor of c-Abl, were detected by western blot analysis with specific antibodies. A representative blot is shown. (C-D) Inhibition of the Abl tyrosine kinase reduces tyrosine phosphorylation of the GRK2-S670A mutant in G2 and restores CDK1 activation. Adaphostin (50µM) or vehicle was added 2h post-release to synchronized HeLa cells expressing GRK2-S670A and maintained for the indicated time points. Tyrosine phosphorylation of immunoprecipitated GRK2 (C) and both phosphorylation and total protein levels of AKT and CDK1 were determined (D). Levels of pY15-CDK1 were maintained in untreated HeLa-A1 cells through G2 (protracted G2) compared to decreasing levels in treated HeLa-A1 cells that normally enter from G2 into mitosis (G2/M progression). pY-GRK2 and p-AKT levels were normalized by total GRK2 and AKT respectively, and represented as fold-change with respect to maximal pY-GRK2 and AKT activation in vehicle-treated HeLa-A1 cells. *, p< 0.05, ** p< 0.01, *** p< 0.001, compared to untreated cells. Blots are representative of 2-3 independent experiments. Dashed lines indicate removal of lanes within the same blot

Both Abl2 and its paralog Abl1, which share more than 90% identity in their SH3-SH2-TK domains, form a unique family of kinases closely related to c-Src (40). Given the recognized role of Abl1 in regulating the G2/M transition (41), we investigated whether this tyrosine kinase interacts with GRK2 in a cellular setting. We observed that both endogenous and overexpressed GRK2 co-immunoprecipitated with overexpressed Abl1 in asynchronously growing HeLa cells (Fig. 3B, left panel). Notably, tyrosine phosphorylation of GRK2 increased concomitantly with Abl1 overexpression but was completely inhibited in the presence of adaphostin, a selective non-ATP inhibitor of Abl tyrosine kinase that blocks the binding site of peptide substrates (Fig. 3B, right panel). Consistent with Abl being responsible for the cell cycle-dependent modification of GRK2, the addition of adaphostin significantly reduced the extent of GRK2-S670A phosphorylation in HeLa cells progressing synchronously through the G2 phase (Fig.3C). Moreover, inhibiting c-Abl activity alleviated the untimely AKT activation triggered by GRK2-S670A, thereby restoring normal CDK1 activation, as indicated by the decrease in its inhibitory phosphorylation levels (pY15-CDK1) (Fig.3D).

These findings suggest that extra levels of improperly stabilized GRK2 facilitate c-Abl-mediated phosphorylation of the protein during the G2 phase, followed by recruitment to p85/PI3K, subsequently leading to AKT activation and delay in cell cycle progression.

### The extent of AKT activation by GRK2 and the delay in the G2 phase in normal cycling cells are subordinated to the transactivation-independent activity of p53

AKT activity can induce various effects on cell cycle progression and survival, depending on factors such as cell type, environmental stress conditions, and the transformed status of the cell. In normal, growth factor-stimulated primary cells, AKT activation promotes permanent cell cycle arrest through p53 activation and inhibition of FOKO3a (42). In contrast, activating AKT in transformed cells attenuates cell cycle arrest independently of p53 (17) and downregulates CHK1 activation (20, 43), the major checkpoint kinase involved in G2/M arrest. Thus, complex interactions between p53 and AKT determine cell fate, involving gene transcription, post-translational protein modification, and lipid modifications (44, 45, 46).

Therefore, the distinct functionality of p53 may influence the contribution of AKT to cell cycle progression and G2 arrest. Hence, we next investigated whether the impact of impaired GRK2 degradation on the G2 phase could involve the participation of p53, either by contributing to unscheduled AKT activation or by delaying G2 transition through other mechanisms. Despite effectively reducing p21 levels, pharmacological treatment of HeLa-K1 cells progressing synchronously into G2 with Pifitrin-α, a specific inhibitor of p53 transcription activity, showed no effect on the high levels of activated AKT (Fig. 4A). Similarly, neither the defective activation of CDK1 nor the impaired upregulation of cyclinB1 and mitotic phosphorylation of histone H3 were affected (Fig. 4A). This suggests that the transactivation activity of p53 is not required for GRK2 accumulation to induce AKT activation or to delay the transition to mitosis in G2.

**Figure 4.**
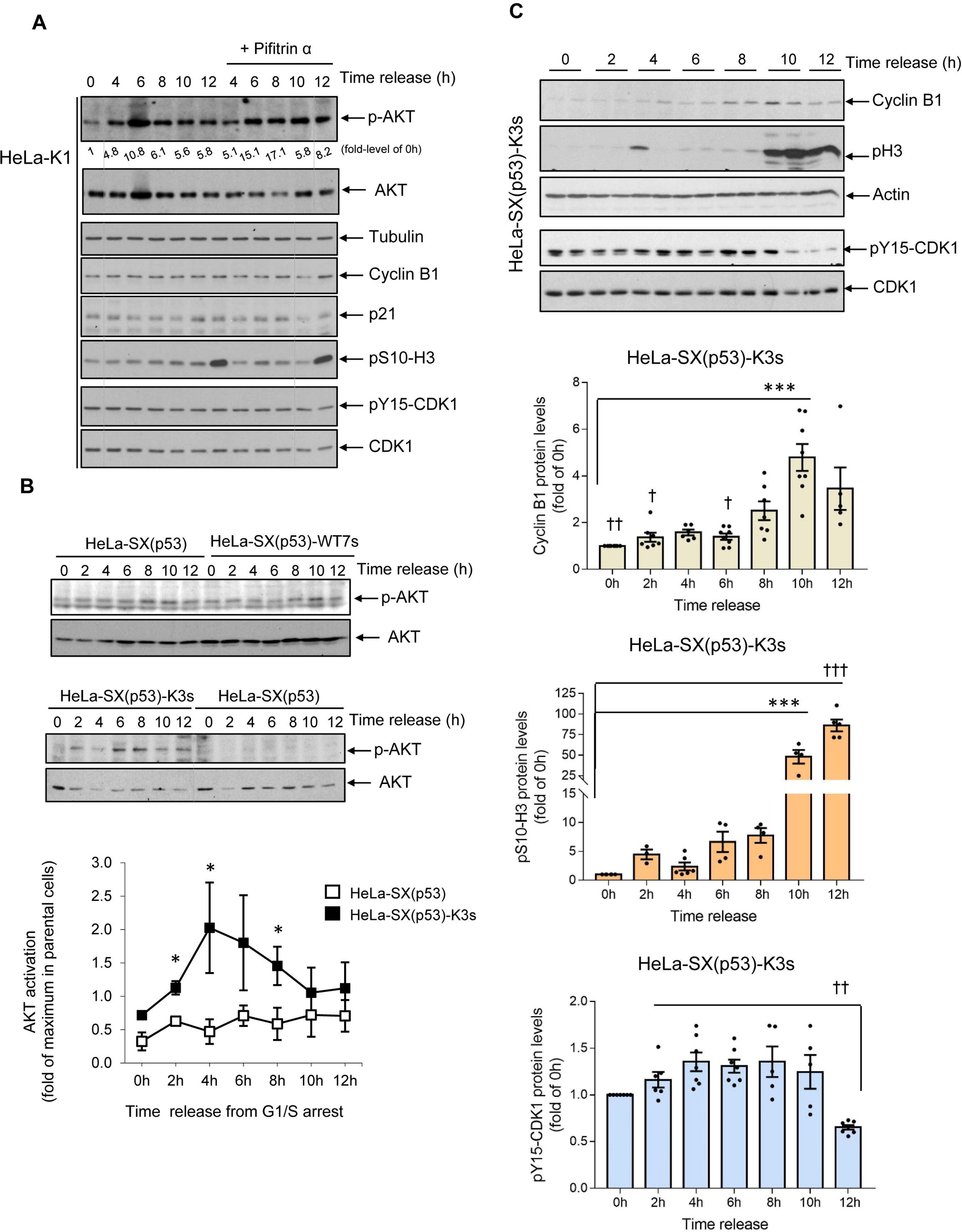
The stimulation of AKT by deficiently downmodulated GRK2 in G2 is fostered by p53 independently of its transactivation activity. (A) The p53 transcriptional activity have no effect on the GRK2-K220R-mediated activation of AKT nor impairment of G2/M transition. Synchronized HeLa-K1 cells were released into G2 in the absence or presence of 40 µM pifithrin-α, a specific inhibitor of p53 transcription activity, for the times indicated. The cell lysates were analyzed by immunoblotting with anti-pS473-AKT (p-AKT), total AKT antibodies, and cell cyle progression markers (Cyclin B1, pS10-H3, pY15-CDK1). The lower levels of basal p21 in pifithrin-treated cells 10-12 h after release from a G1/S arrest confirm the effective inhibition of p53-dependent transactivation. Similar results were obtained in 2 separate experiments. (B-C) In the absence of p53 expression, the impaired downregulation of GRK2 in cells progressing through G2 leads to a less prominent activation of AKT without protracting effects on G2/M transition. (B) Control parental cells with silenced p53 (HeLa-SX(p53)) and cells stably overexpressing GRK2-wt (HeLa-SX(p53)-WT7s) or GRK2-K220R (HeLa-SX(p53)-K3s) were synchronized at the G1/S boundary for subsequent release into cell cycle, and cellular lysates processed for the immunodetection of p-AKT and total AKT protein at the indicated times. Normalized p-AKT levels were represented as fold-change with respect to maximal AKT activation in control parental HeLa-SX(p53) cells. Data are mean ± SEM of 2-3 independent experiments. *, p< 0.05, ** p< 0.01, *** p< 0.001, compared to parental cells at the corresponding times. (C) HeLa-SX(p53)-K3s cells timely progress through G2 and mitosis. Cells were synchronized at the G1/S transition and released into the cell cycle for the time indicated, and cellular lysates were immunoblotted with anti-pS10-H3, anti-cyclin B, and anti-Y15-CDK1 antibodies. After stripping, blots were probed with anti-actin and anti-CDK1 antibodies for control loading. Data are mean ± S.E.M of 3-8 independent experiments. *** p< 0. 001 and † p< 0.05, †† p< 0.01, ††† p< 0.01, one-way Anova test when compared to 10 h and 12 h post-release, respectively. Representative blots are shown. Dashed lines indicate removal of lanes within the same blot.

To further investigate the role of p53, we utilized HeLa cells in which p53 was permanently silenced using a specific shRNA sequence (referred to as HeLa SilenciX® p53 or HeLa-SX(p53) hereinafter). This engineered cell line exhibited no detectable p53 protein expression, in contrast to the reduced levels observed in parental HeLa cells. This difference was evident under subtle conditions of p53 accumulation, such as during routine progression through late G2 phase and mitosis, or under strong induction of the p53 protein, as seen during cell cycle arrest caused by DNA damage-induced checkpoint activation in the G2 phase (Suppl.Fig.S4A).

We generated stable cells lines overexpressing either GRK2-wt (designated as HeLa-SX(p53)-WT7) or a degradation- and catalytic-deficient mutant of GRK2 (HeLa-SX(p53)-K3), achieving expression levels comparable to those obtained in stable HeLa-WT5 and HeLa-A1 or HeLa-K1 cells (Suppl.Fig.S4A). Further confirming the suitability of the HeLa-SX (p53) cellular model, the absence of p53 had no impact on the cell cycle-dependent decay of endogenous GRK2 (Suppl.Fig.S4C, upper panel) or overexpressed wild-type GRK2 during the G2/M transition (Suppl.Fig.S4C, central panel). Additionally, it did not affect the stabilization of GRK2 protein in response to G2 checkpoint activation (upper and central panels) or the impaired downregulation of GRK2-K220R protein in G2 (Suppl.Fig.S4C, bottom panel).

The observed effect of GRK2 on the AKT pattern in HeLa cells progressing synchronously through G2 was roughly replicated in the absence of p53. Thus, the levels of pAKT in HeLa-SX(p53) cells were unaffected by the overexpression of wt-GRK2 in HeLa-SX(p53)-WT7 cells, while they appeared to increase in HeLa-SX(p53)-K3 cells (Fig. 4B). However, the AKT activation observed in the latter cells (approximately 2-fold versus parental cells) was far less pronounced compared to the HeLa-K1 counterpart (approximately 20-fold). This suggest that multiple pathways may converge in the AKT activation triggered by the GRK2-K220R mutant, with some being independent of p53 activity and others dependent on the transactivation-independent properties of the p53 protein.

Surprisingly, in HeLa-SX(p53) cells, the expression of GRK2-K220R, despite increasing the activation of AKT, had no effect on delaying cell cycle progression at the G2/M transition. Instead, a clear and timely upregulation of Cyclin B1, activation of CDK1, and mitotic phosphorylation of Histone 3 were observed (Fig. 4C), which contrasts with the protracted effect induced by this mutant in regular HeLa cells (Suppl.Fig.S1C). Overall, these results suggest that in the context of the latter cells, the enforced GRK2/p85-PI3K activation collaborates with a p53-mediated, transactivation-independent activation of AKT (47), thus leading to a pronounced FOXO3a phosphorylation by AKT. This modification inhibits FOXO3a’s transcriptional activity in late G2, consequently delaying the onset of mitosis.

### The GRK2-mediated activation of AKT is a p53-independent component of the DNA damage-induced G2-checkpoint

Given that stimulation of AKT has also been observed in response to DNA damage through complex mechanisms involving ATM, ATR, and DNA-PK (48), we aimed to determine whether the natural stabilization of GRK2 during the G2 checkpoint (22) might contribute to the over-activation of AKT in this setting and the resulting impact on cell cycle arrest under genotoxic conditions.

We found that in doxorubicin-treated parental HeLa cells progressing synchronously through the cell cycle, the accumulation of GRK2 at the G2 phase coincided with the activation of AKT and the inhibition of CDK1 (Fig. 5A). Similar to the regulation of basal AKT in stable HeLa-GRK2_436-689_ cells (where endogenous GRK2-WT is enforced to be stabilized during G2 phase), the higher the accumulation of GRK2 protein in response to doxorubicin, the greater the level of AKT activation in G2 checkpoint-arrested cells, as observed in HeLa-WT5 cells compared to parental cells (Fig. 5B). Additionally, in HeLa-A1 cells, the doxorubicin-induced stimulation of AKT was even more pronounced, consistent with the intrinsic overactivation of basal AKT in these cells during the G2 phase (Fig. 5B). In both HeLa-WT5 and HeLa-A1 cells, the addition of LY294002 completely eliminated the activation of AKT by doxorubicin (Fig. 5C), indicating that stabilized GRK2 in G2 checkpoint-arrested cells also regulates AKT through PI3K activation. Furthermore, even though the levels of p-AKT were lower in doxorubicin-treated HeLa-SX(p53) cells compared to normal HeLa cells (Suppl.Fig.S5A), similar regulation by GRK2 was observed in stress-induced AKT. Activation of AKT increased in both parental HeLa-SX(p53) cells and in those stably overexpressing GRK2-wt or catalytic- and degradation-deficient mutant proteins (HeLa-SX(p53)-K3s) when challenged with doxorubicin (Suppl.Fig.S5B-C), paralleling the extent of stabilization and accumulation of GRK2 protein (Suppl.Fig.S4C).

**Figure 5.**
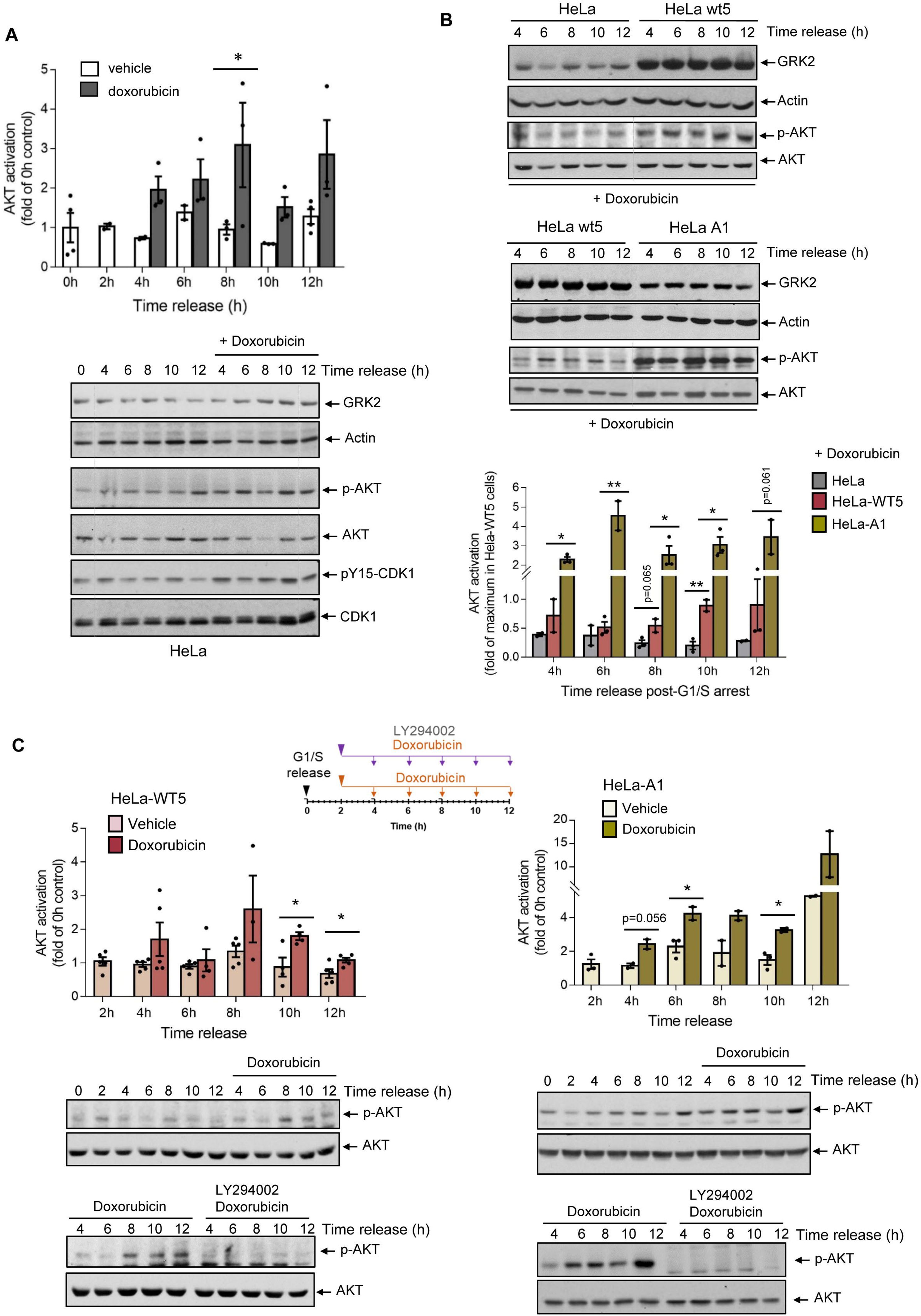
Induced accumulation of GRK2 by DNA damage contributes to PI3K-dependent activation of AKT in G2 arrested cells. (A, B) Doxorubicin-triggered AKT activation in the G2-checkpoint response of HeLa cells correlates with accumulated GRK2. (A) Two hours after release from the block at G1/S, synchronized HeLa cells were exposed to doxorubicin (1 μM) or vehicle for the indicated times, and the protein levels of GRK2, actin, total and phospho-CDK1 (pY15-CDK1), and total and phospho-AKT (p-AKT) were analyzed with specific antibodies. Levels of p-AKT were normalized by total AKT and represented as fold-change with respect to 0h AKT activation in HeLa cells. Data are mean ± SEM of 3-4 independent experiments. *, p< 0.05, compared to vehicle-treated cells at each time. (B) Parental HeLa cells and cells expressing levels of extra GRK2-wt (HeLa-WT5) or GRK2-S670A (HeLa-A1) were synchronized and treated with doxorubicin as above, and lysates collected at the indicated times to analyze GRK2 protein levels and phosphorylation status of AKT. Normalized p-AKT levels were represented as fold-change with respect to maximal AKT activation in HeLa-WT5 cells. Data are mean ± SEM of 2-3 independent experiments. *, p< 0.05, ** p< 0.01, compared to GRK2-wt overexpressing cells. (C) Synchronized HeLa-WT5 and HeLa-A1 cells were exposed to vehicle or doxorubicin as above in the presence or absence of PI3K inhibitor LY294002 (10µM). Total and phospho-AKT (p-AKT) levels were analyzed with specific antibodies, and normalized p-AKT represented as fold-change with respect to AKT activation at 0h released cells. Data are mean ± SEM of 2-5 independent experiments. *, p< 0.05, compared to vehicle-treated cells at each time. Representative blots are shown. Dashed lines indicate removal of lanes within the same blot.

Overall, these findings indicate that the GRK2-mediated and PI3K-dependent activation of AKT is an inherent signaling pathway within the G2 checkpoint response to DNA damage, regardless of the presence or absence of p53.

### GRK2-mediated activation of AKT reinforces the initial cell cycle arrest in the G2-checkpoint while facilitating the abrogation of sustained arrest, dependent on p53

While DNA damage triggers AKT activation in various cellular settings (17, 18, 19, 20, 48), conflicting results have been reported on the effects of AKT on cell cycle progression when the G2 checkpoint is activated. These results range from contributing to cell cycle arrest to overcoming the pause imposed by the G2 checkpoint. Therefore, we investigated the functional effect of GRK2-mediated AKT activation on the dynamics of markers of the G2/M transition that are inactivated by the G2 checkpoint. Correlating with the higher levels of p-AKT (Fig 5B), doxorubicin-induced downmodulation of Cyclin B1 and pS10-H3 protein was more intense in HeLa-WT5 cells compared to parental cells (Fig. 6A). Meanwhile, in the absence of damage, both cell lines progressed similarly from G2 into mitosis. Notably, the addition of LY294002 relieved AKT over-activation by doxorubicin in HeLa-WT5 cells, leading to a weaker induction of cell cycle arrest, as evidenced by partial recovery of cyclin B1 and pS10-H3 levels and activation of CDK1 (Fig. 6B).

**Figure 6.**
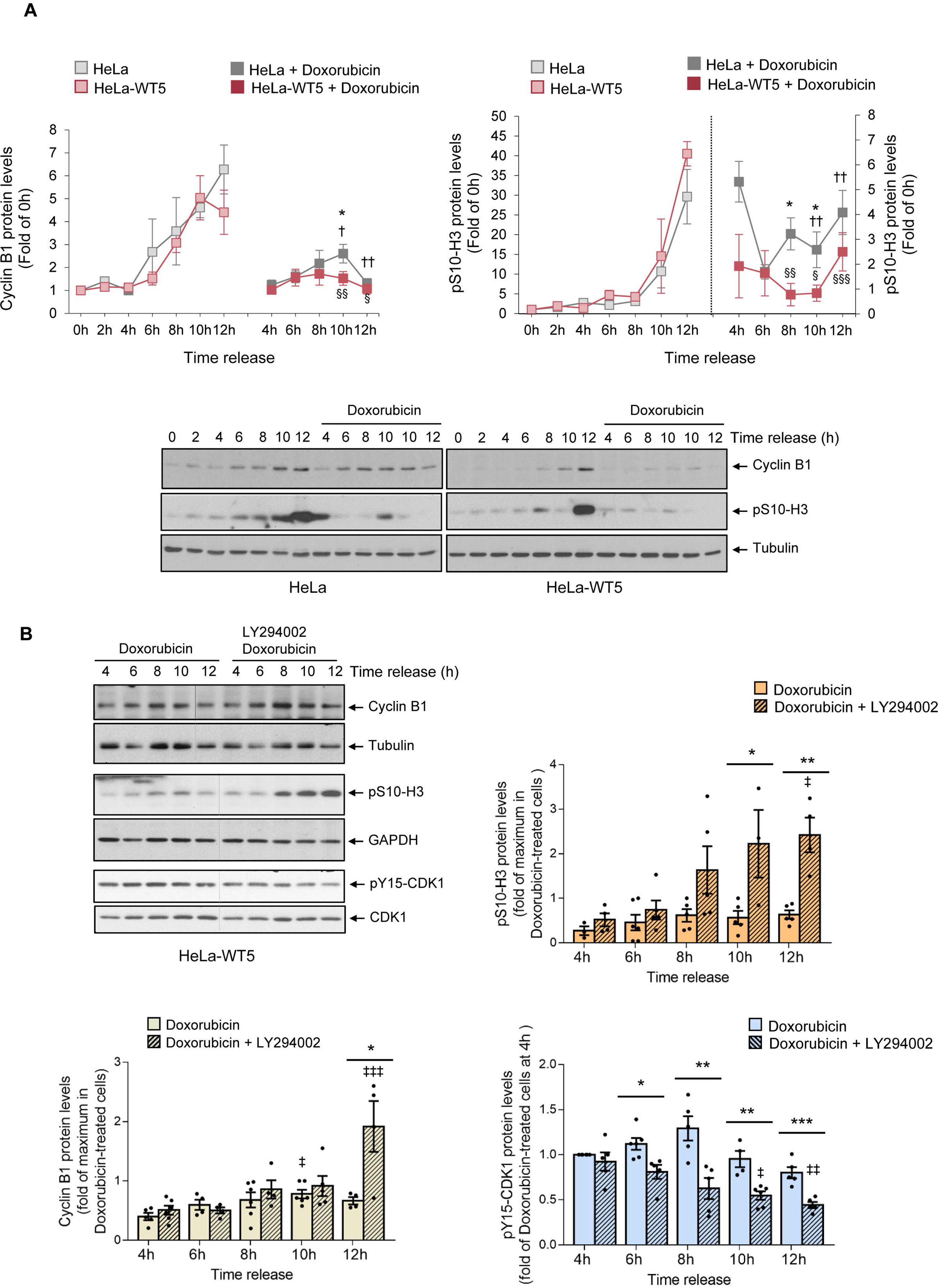
Doxorubicin-triggered GRK2 stabilization strengthens the induction of G2 checkpoint arrest with the involvement of PI3K. (A) GRK2 accumulation by doxorubicin correlates with the extent of inhibition of mitotic drivers (cyclin B1) and markers (pS10-H3) in the G2 checkpoint response. Synchronized HeLa cells with or without extra wild-type GRK2 (HeLa-WT5) were exposed to doxorubicin (1 µM) or vehicle two hours post-release from a G1/S block for the indicated times, and levels of Cyclin B1, phosphorylated H3 (pS10-H3), and tubulin proteins were immunodetected with specific antibodies. Cyclin B1 and pS10-H3 densitometry values were corrected for tubulin expression and represented as fold-change with respect to 0h release. Data are mean ± SEM of 2-6 independent experiments. * p< 0.05, when compared doxorubicin-treated parental with HeLa-WT5 at the corresponding time. † p< 0.05, †† p< 0.01, when compared doxorubicin-treated with untreated HeLa cells and § p< 0.05, §§ p< 0.01, §§§ p< 0.001 for comparison between doxorubicin-treated and untreated HeLa-WT5 cells at the corresponding time. (B) PI3K inhibition relieves the cell cycle arresting effects of stabilized GRK2 protein in the G2 checkpoint response. Synchronized HeLa-WT5 cells as above were exposed to vehicle or doxorubicin in the presence or absence of PI3K inhibitor LY294002 (10µM). Cellular lysates were subjected to immunoblot analysis to determine expression levels of Cyclin B1, pS10-H3 and total CDK1 and pY15-CDK1. Levels of normalized pY15-CDK1 by CDK1 were represented as fold-change with respect to 4h doxorubicin-treated cells, and levels of Cyclin B1 and pS10-H3, respectively corrected by tubulin and GAPDH content, with respect to the maximum value in doxorubicin-treated cells. Data are mean ± SEM of 3-6 independent experiments. * p< 0.05, ** p< 0.01, *** p< 0. 001, when compared LY294002-inhibited with non-inhibited cells at the indicated time points. ‡ p< 0.05, ‡‡p< 0.01, ‡‡‡ p< 0. 001, one-way Anova test when compared doxorubicin plus LY294002 treated cells at different times with respect to 4h. In all panels representative blots are shown. Dashed lines indicate removal of lanes within the same blot

The p53 protein can play a relevant role in blocking the G2/M transition by inhibiting CDK1 through different mechanisms (14), and in HeLa cells, p53 induction by DNA damage contributes to the full activation of AKT at the G2 checkpoint (Suppl.Fig.S4A). Hence, we addressed whether the GRK2-mediated p53-independent component of AKT itself could enhance cell cycle arrest in cells challenged with doxorubicin upon release of a G1/S block. The down-modulation of cyclin B1 and pS10-H3 protein levels caused by doxorubicin was higher in HeLa-SX(p53)-WT7s and HeLa-SX(p53)-K3s cells compared to parental HeLa-SX(p53) cells (Suppl.Fig.S6A). Similar attenuation of CDK1 activation was observed, with clearly lower levels in HeLa-SX(p53)-K3s cells and a trend in HeLa-SX(p53)-WT7s cells (Suppl.Fig.S6A). These observations strongly suggest that AKT activation through the stabilization of GRK2 contributes to reinforcing cell cycle arrest independently of p53 in the first hours of genotoxic damage.

It is widely accepted that the initiation of G2 arrest following DNA damage does not rely on the function of p53, in line with the similar effectiveness of doxorubicin-induced changes in Cyclin B1 or pS10-H3 protein levels between parental HeLa and HeLa-SX(p53) cells (Fig. 6A and Suppl.Fig.S6A). Nevertheless, the maintenance of G2 checkpoint arrest is known to require the involvement of p53 (15, 16, 49). On the other hand, chronic activation of AKT has been reported to overcome long-term arrest by the G2/M checkpoint independently of p53 (17, 19). Therefore, we investigated whether the stabilization of GRK2 protein levels by DNA damage could alter the cell’s ability to maintain G2 checkpoint arrest over time in different p53 contexts. To examine this, asynchronously growing cells were treated with doxorubicin for 24 h and cell cycle profile was determined by the measurement of cellular DNA content using flow cytometry. Normal cells with competent checkpoints are arrested in either the G1 phase or G2 phase, but tumor cells usually cannot arrest in the G1 phase following DNA damage. In this context, HeLa cells, characterized by a weakened p53 response, were primarily arrested at the G2/M transition after 24 h of doxorubicin treatment in a dose-dependent manner (Fig 7A). Neither the complete elimination of p53 nor the overexpression of wild-type and catalytically inactive forms of GRK2 in HeLa cells altered the point at which cell cycle arrests in the presence of doxorubicin (Fig 7A, B). However, the proportion of G2 checkpoint-arrested cells differs in both parental and stably transfected HeLa and HeLa-SX(p53) cell lines. In the presence of p53, the maintenance of the G2/M arrest was overridden by increased levels of wild-type GRK2 protein at the lowest dose of doxorubicin (0.05 µM), and by kinase-dead protein at both low and high doses (Fig 7A). This phenomenon was not observed in HeLa-SX (p53) cells (Fig 7B), pointing to an inverse correlation between G2 arrest maintenance and GRK2-mediated activation of AKT only in p53-positive cells. Moreover, this less stable G2 arrest involves the successful exit from G2/M arrest and re-entry into the cell cycle as confirmed in doxorubicin-treated HeLa-WT5 and HeLa-K1 cells plated at low density for long-term colony formation. In comparison to the parental cells and their counterparts in the HeLa-SX(p53) background, these cells exhibited a markedly increased survival fraction (Fig 7C, D). Interestingly, the clonogenic survival of HeLa-SX(p53) cells was only modestly improved in the presence of GRK2-K220R protein (Fig.7 D), a mutant leading to a more pronounced impact on AKT activation (Suppl. Fig. S5D). Taken together, these results provide substantial evidence that the convergence of GRK2/p85 and p53 signaling plays a pivotal role in regulating AKT, which, in turn, determines the fate of cellular responses to the G2 checkpoint, ultimately influencing whether genotoxic-treated cells undergo cell death or survival.

**Figure 7.**
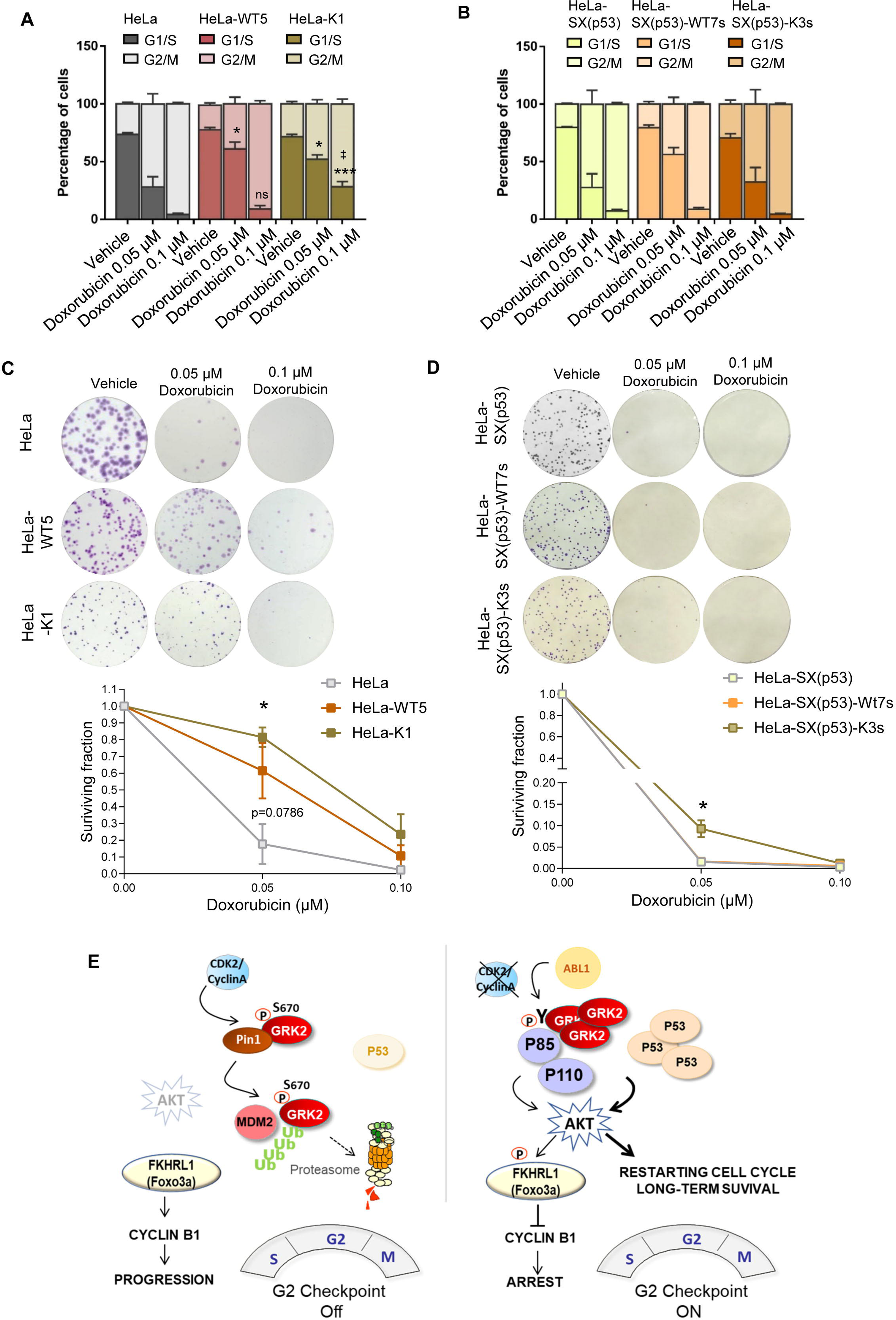
The genotoxic-induced accumulation of GRK2 facilitates the overriding of long-term cell cycle arrest and survival in a p53-dependent manner. (A, B) In HeLa cells exposed to 24h of doxorubicin, but not in HeLa-SX(p53) cells, extra levels of GRK2 lead to a significant reduction in the number of G2/M arrested cells. Asynchronously growing HeLa (A) and p53-silenced HeLa-SX(p53) (B) cells, with or without overexpressing wild-type GRK2 or GRK2-K220R mutant, were treated with 0.1 or 0.05 µM doxorubicin for 24h and harvested for flow cytometric analysis of DNA content as detailed in Methods. The percentage of cells in different stages of the cell cycle was quantified and represented. Data are mean ± SEM of six (A) and three (B) independent experiments. * p< 0.05, *** p< 0.001, when compared to the percentage of GRK2-overexpressing cells at G1/S with parental cells, and †† p< 0.01 when comparing GRK2-wt versus GRK2-K220R overexpressing cells. (C, D) Extra GRK2 levels increase long-term cellular survival upon doxorubicin treatment only in the presence of p53. The same cells as in panels A, B were also seeded into 6-well plates and cultured for 10 days after 24h of doxorubicin exposure as detailed in Methods. Colonies were visualized with Crystal violet. Representative images are shown. Clonogenic survival was quantified as described in Methods, and data plotted as a fraction relative to untreated cells. Data are mean ± SEM of four (C) and three (D) independent experiments performed in duplicate. * p< 0.05, *** p< 0.001, when compared to doxorubicin-treated parental HeLa and HeLa-SX(p53) cells. (E) Scheme of GRK2 regulation during G2 progression and molecular model for GRK2-mediated activation of AKT at the G2 checkpoint. GRK2 is intrinsically degraded in cycling cells by a pathway involving CDK2/cyclinA-mediated phosphorylation and functional interaction with Pin1. This enables recognition and ubiquitination by MDM2, restraining AKT activation and ensuring timely cell cycle progression at G2/M (22, 24). At the G2 checkpoint, GRK2 protein stabilizes concurrently with p53 upregulation and AKT activation. Accumulated GRK2 at G2 is phosphorylated by Abl1, allowing direct association with the p85 regulatory subunit of PI3K through N-terminal phosphotyrosine residues and C-terminal proline box interacting motifs (see text for details). This GRK2-dependent, PI3K-mediated stimulation of AKT causes a pro-arresting inhibition of Foxo3a-dependent Cyclin B increase. The interplay of the Abl1/GRK2/p85-based class 1A PI3K signalosome with p53-mediated transactivation-independent mechanisms sustains AKT activation, aiding cells in resuming cell cycle and surviving after massive genotoxic damage.

## DISCUSSION

We uncover that DNA-damage-mediated upregulation of GRK2 plays a pivotal role in halting cell cycling at the G2 checkpoint in the absence of p53. Conversely, in a p53-dependent manner, GRK2 facilitates the exit from G2 arrest, promoting long-term survival. Both effects rely on the increased activation of AKT and the tilting of p53’s role from apoptosis to survival in the context of genotoxic damage. Moreover, GRK2-mediated sustained activation of the PI3K/AKT pathway in G2 induces hyper-phosphorylation of the forkhead transcription factor FKHRL1 (FOXO3a), disrupts Cyclin B1 accumulation, and alters CDK1 activation. This, in turn, results in a significant delay in the G2/M transition and overall cell cycle progression. Our findings also underscore the contribution of p53 to chemotherapeutic resistance, highlighting the involvement of GRK2 in AKT activation.

In normal cycling cells, GRK2 is downregulated during G2 progression due to phosphorylation by CDK2 and Pin1 interaction, followed by Mdm2 ubiquitination and degradation (22, 24). We now demonstrate that this process is key to prevent unscheduled PI3K-dependent activation of AKT, thereby ensuring the G2/M transition. Disruption of this physiological regulatory interplay, by expression of GRK2 mutants resistant to CDK2- and Pin1-dependent degradation, or inhibition of endogenous GRK2 downmodulation using CDK2 inhibitors, results in the direct association of accumulated GRK2 with the p85 PI3K regulatory subunit and subsequent activation of the PI3K/AKT signalling pathway (Fig. 7E). Our results further indicate that an “excess” of GRK2 during G2 promotes its phosphorylation at tyrosine residues by the cell cycle regulatory kinase c-Abl, enhancing its association with the p85 PI3K subunit. This mechanism, crucial for the arresting effect of excess GRK2 in G2, can be mitigated by blocking tyrosine phosphorylation of GRK2, leading to attenuated pAKT levels and the resumption of cell cycle progression.

We have demonstrated a direct binding capability of GRK2 to the p85 regulatory subunit of PI3K, in contrast to the previously observed interaction with the p110 catalytic subunit (35). While GRK2 interaction with the p110 subunit has been established for PI3Kγ, and suggested for other PI3Ks based on GRK2 recruitment to p110γ via the common PIK domain, we find that both the RH domain and the 467-474 proline-motif of GRK2 are involved in p85 binding. This contrasts with prior reports indicating that the C-terminal portion of GRK2, excluding the proline-box, binds to the PI3K catalytic subunit (50). Notably, our observation that a PIK construct does not disrupt the GRK2-p85 interaction suggests diverse mechanisms for different PI3Ks in recruiting GRK2. Thus, Class IA PI3Ks appears to preferentially display a p85-mediated association, while class IB exhibits a p110 and PIK-mediated association.

The basal inhibition of p110 in the PI3K p85/p110 complex is relieved through the interaction of the p85 SH2 domains with Tyr-phosphorylated RTK and adaptor proteins (51, 52, 53). Additionally, cytosolic Tyr kinases, such as Src family kinase members, can activate PI3K through a distinct mechanism involving their SH3 domains and the p85 Pro-rich regions (54). Our results show that GRK2 association with p85 involves such SH2 and SH3 domains, is dynamically modulated by phosphorylation, and potentially contributes to PI3K activation. While phosphorylation of GRK2 at Ser670 does not directly modulates p85 binding, it critically influences the levels of GRK2 available for association. Phosphorylation at tyrosine residues 86 and 92 significantly enhances GRK2/p85 interaction. c-Abl-mediated tyrosine-phosphorylation of GRK2 during G2 increases its association with p85, potentially leading to PI3K activation akin to the activation mode of adaptor proteins (51). The binding of GRK2 to p85 also involves the Pro-rich region of GRK2 and the SH3 domain of p85, similar to the activating interaction between PI3K-p85 and c-Src. However, it remains plausible that GRK2 may form a complex with free p85 subunits in the absence of p110, indirectly impacting on PI3P levels. For instance, the proline-rich motif-mediated interaction of PRR11 with the p85α subunit suppresses p85 homodimerization and competes with p85/110 heterodimers for binding to RTKs (55). The estrogen receptor also activates PI3K/AKT through an interaction with p85α via regions other than the SH2/SH3 domains (56). On the other hand, apart from forming complexes with p110, there is a pool of p85 associated with PTEN through the N-terminal SH3 domain, enhancing PTEN lipid phosphatase activity (57). Therefore, GRK2 could increase the PIP3 pool by competing with PTEN for binding to p85, indirectly activating the PI3K/AKT axis. Despite the potential broader implications of the novel GRK2/p85 interaction in cellular contexts where kinase upregulation leads to AKT activation, such as migration or mitogenic signaling (21, 58, 59), the role of GRK2 in unleashing p85 inhibitory control on PI3K is intrinsic to the cell cycle under genotoxic stress. Doxorubicin treatment disrupts the downmodulation of GRK2, correlating with sustained AKT activation and stronger cell cycle arrest induction.

Non-canonical nuclear AKT activation may occur following DNA damage involving cell cycle-related kinases. Genotoxic stress activates the PIKK family members ATM, ATR and DNA-PK, along with a complex cascade of kinases impacting AKT regulation through mechanisms not fully understood (48). For instance, ATM directly binds to AKT but does not phosphorylate it, suggesting intermediate proteins in its activation (60), while DNA-PK directly phosphorylates and activates AKT (61). Although under certain circumstances active DNA-PK may play a role in G2 checkpoint (62), a role in GRK2-mediated activation of AKT and concomitant phosphorylation of FOXO3a is unlikely due to its entirely dependency on PI3K activity and the reported inability of DNA-PK-activated AKT to phosphorylate FOXO3a (63).

On the other hand, CDK2/Cyclin A is known to periodically phosphorylates AKT at its carboxyl terminus (S477/T479) during the S/G2 transition and early G2 phase (64). This modification may facilitate or compensate for canonical S743 phosphorylation in the basal cell cycle, as a threshold activity of AKT is necessary for progression (30). In normal cycling cells, this activation mechanism aligns with the CDK2-Cyclin A1-induced GRK2 downmodulation (22), stressing its relevance in the maintenance of appropriate levels of AKT activity. Though CDK2-Cyclin A1 was reported to promote the G2/M checkpoint in some contexts and phosphorylates AKT (65), its contribution in our model is also unlikely since CDK2 is inactivated in response to DNA damage in p53-expressing cells and its activity is dispensable for G2 arrest by doxorubicin (66). Indeed, inactivation of Cdk2 as part of the DDR results in the stabilization and accumulation of GRK2, strongly delaying G2 progression upon DNA damage (22, 67). Overall, GRK2 scaffold-mediated regulation of p85 activity is the most likely mechanism by which doxorubicin-induced GRK2 or the accumulation of degradation-compromised mutants stimulates the PI3K/AKT axis.

c-Abl1-mediated tyrosine phosphorylation of GRK2 and subsequent scaffolding-dependent activation of PI3K might constitute a central process ensuring an efficient G2 checkpoint cell cycle arrest. Upregulation of c-Abl1, known to occur in response to doxorubicin in U2OS and HeLa cells, is associated with pro-arresting and anti-apoptotic effects in the G2 checkpoint (68). In these cellular settings, inhibiting c-Abl1 significantly reduced DNA damage-induced G2 arrest, even in the presence of substantially induced p53. c-Abl1 activity regulates the DNA damage checkpoint response through both p53-dependent and independent mechanisms, modulating cell fate decision to survival, senescence, or apoptosis (68, 69). Additionally, hyperactivation of c-Abl1 in tumoral contexts, as seen with the oncogenic BCR-ABL1 chimeric protein fusion, inhibits apoptosis and prolongs G2/M arrest, resulting in genotoxic resistance (70). Therefore, it is feasible that GRK2-dependent stimulation of AKT might play a role in mediating the genotoxic resistance triggered by c-Abl1.

Cancer cells’ resistance to apoptosis following DNA damage is linked to the activation of survival pathways, coupled with mechanisms that potently halt the cell cycle. The robustness and duration of G2 checkpoint activation directly correlates with the cellular resistance to genotoxic agents (1, 11). There is evidence of AKT exhibiting a dual role in G2/M checkpoint regulation depending on cell type and signaling context (48). In epithelial cells, PI3K/AKT inhibition induces G2 arrest, while constitutively active AKT allows cells to bypass genotoxic-induced G2 checkpoint and progress into mitosis (17, 20). Conversely, in hematopoietic cells, PI3K/AKT signalling strengthens G2 checkpoint activation in response to DNA-damaging Topoisomerase II inhibitors (18). Other studies have demonstrated AKT overriding stable G2 arrest after γ-irradiation (19).

Our findings reveal that genotoxic-induced upregulation of GRK2 facilitates AKT activation, resulting in a more potent early G2 checkpoint arrest and easier cell cycle resumption at later times for long-term survival. AKT, with numerous substrates influencing G1/S and G2/M transitions in unperturbed cycling cells and checkpoint responses to DNA damage (48), is well-established for its positive role in cell growth, facilitating mitogenic entry into G1, promoting transition to mitosis, and preventing apoptosis. AKT achieves transition to mitosis by directly phosphorylating Cdc25 (71), inactivating negative regulators of CDK1/Cyclin B1 (72, 73), and phosphorylating Mdm2, leading to p53 attenuation (74). However, emerging evidence suggest anti-proliferative and pro-death effects of AKT activity through the p53/p21-dependent pathway (42, 48, 75). Moreover, transient AKT activation is essential for normal cell cycle progression (30); however, sustained activation leads to G2/M arrest and apoptosis due to CDK2 mislocalization (76). AKT phosphorylates and relocates FOXO3a to the cytosol through the interaction with 14-3-3 proteins, resulting in mitotic arrest (33, 77). Our findings offer additional insights, highlighting the necessity of p53 for the pro-arresting effect of hyper-activated of AKT and FOXO3a phosphorylation. Considering that 14-3-3 is a transcriptional target of p53 (78), it is possible that in Hela-p53 null cells, the redistribution of FOXO3a may be inefficient. Various more intricate scenarios may exist, given the intricate crosstalk between FOXO3a and p53, displaying cooperation or antagonism depending on the cellular context. FOXO3a physically interacts with p53, regulating both transcriptional and transactivation-independent functions of p53 (79). Additionally, these factors share transcriptional targets that induce cell cycle arrest, such as GADD45A (80), so their potential convergence in promoting cell cycle arrest cannot be ruled out.

Intriguingly, in cells lacking p53 expression, we observe diminished levels of AKT activation mediated by GRK2 in both unstressed cells and in those timely induced at the G2 checkpoint. This correlation aligns with attenuated pro-arrest and survival responses. This observation suggests that if the activation of AKT does not surpass a certain threshold, there may not be sufficient active AKT to functionally override a significant number of FOXO3a molecules and halt the cell cycle. It also implies that the p53 protein might collaborate in AKT activation during genotoxic stress in the G2 checkpoint, particularly in the presence of degradation-defective GRK2 mutants during G2 progression. Notably, AKT has been identified as a potential p53 partner, and recent findings indicate that p53 assembles a PI3K-AKT pathway that regulates AKT activation independently of the canonical membrane-localized activation of PI3K (47). A nuclear phosphoinositide kinase-p53 complex has been identified in response to stress, producing PtdIns (4,5)P2 that remains associated with p53, thereby regulating its interaction with other proteins (81) and serving as a substrate for nuclear PI3K (47). In this context, is tempting to suggest that GRK2 may play a role in facilitating the function of p53 in recruiting AKT-activating kinases. Analogous to how p85 SH2 domains bind to phosphorylated RTKs and localize p110 to the plasma membrane where its substrate resides, non-membrane locations could be achieved through the interaction of p85 with phospho-tyrosine GRK2. Indeed, although the primary location of the PI3K complex is in the cytosol, nuclear activity has also been described under certain stimuli (47). Furthermore, GRK2 has been reported in complex with p53 (82). Therefore, the interaction of GRK2 with p85 could potentially serve to dock the PI3K enzyme in the signalosome of PtdIns (4,5)-associated p53.

The resistance of tumor cells to chemotherapy is often attributed to the absence or impaired activation of p53. Strategies involving the upregulation of p53 expression through gene or small drug-based therapies are suggested to overcome this resistance, expecting more efficient killing of tumors retaining wild-type p53. However, such tumors don’t consistently show improved chemotherapy outcomes, as p53, besides its well-known role as a pro-apoptotic tumor suppressor, functions as an active pro-survival factor (83). Our findings support the idea that the absence of p53 sensitizes cancer cells to death upon DNA damage rather than promoting survival. This unveils a protective mechanism involving the interplay of GRK2/p85-PI3K and p53 to activate AKT, potentially diverting p53’s activity toward cell-cycle arrest and repair instead of apoptosis.

Several factors influence p53’s life and death decisions, including protein concentration, subcellular localization, or posttranslational modifications (9). Additionally, the interplay of p53 with other tumor suppressors and oncogenes, such as the PTEN-PI3K-AKT-MDM2-p53 network (44, 45), plays a crucial role in balancing cell survival and death signaling (84). Within this network, genotoxic inhibition of Mdm2 stabilizes p53 which in turn transactivates Mdm2, delaying apoptosis. On the other hand, PI3K activates AKT, promoting MDM2 nuclear translocation to reduce p53 levels and p53-mediated repression of PI3K (46). PTEN, by inhibiting PI3K/AKT, protects p53 and facilitates PTEN transactivation (85). In this context, GRK2 contributes to raising the apoptotic threshold in stress conditions through AKT activation and networking with p53, reinforcing the idea that p53’s pro-apoptotic role depends on AKT inhibition (84). Consequently, silencing GRK2 expression sensitizes breast cancer cells to the apoptotic effects of genotoxic drugs (21).

Our findings unveil a pivotal role for a scaffold function of GRK2 in the cell cycle regulation of AKT which is engaged upon genotoxic activation of the G2 checkpoint. This mechanism may hold significance for most tumor cells, potentially impacting the pathophysiology of cancer. The GRK2 gene undergoes amplification in specific human cancers, such as breast tumors and esophageal cancer, while protein upregulation is frequently observed in luminal breast tumors or pancreas cancer (86). Increased GRK2 levels influence key tumoral features such as sustained mitogenic signalling (21) or sub-optimal activation of p53 (22, 82). Paradoxically, this latter effect has the potential to better promote survival compared to non-activation (87). Therefore, elevated GRK2 in breast cancer patients may facilitate a pro-survival cellular state (low p53 and high AKT levels), whereas GRK2 downmodulation promotes a pro-apoptotic state (high p53 and low AKT levels). Our results also suggest that inhibiting GRK2 catalytic activity might be less efficient in promoting apoptosis than interfering with gene expression. The combined use of GRK2 kinase silencers and DNA-damaging drugs in chemotherapy against tumors retaining wild-type p53 could be an interesting avenue to explore.

## AUTHOR CONTRIBUTIONS

Conceptualization: P.P; Methodology: V.R, T.G-M., A.A., V.L.; Investigation: V.R, T.G-M., A.A., V.L.; Formal Analysis: V.R, V.L, P.P.; Visualization: V.R, T.G-M., V.L, P.P.; Resources: V.L., P.P.; Writing – Original Draft: P.P; Writing – Review & Editing: P.P., F.M.; Funding acquisition: P.P., F.M.; Supervision: P.P.

## Material and Methods

### Cell culture and cellular transfections

HeLa cells (American Type Culture Collection, Manassas, VA, USA) and HeLa SilenciX® p53 (referred as HeLa-SX(p53) (RRID:CVCL KT87, Tebu-Bio Cat no. 00301-00024) were maintained in Dulbecco’s modified Eagle’s medium (DMEM) supplemented with 10% (v/v) fetal bovine serum (FBS, Sigma-Aldrich, St. Louis, MO, USA), at 37°C in a humidified 5% CO2 atmosphere. HeLa cells (70-80% confluent monolayers in 60- or 100-mm dishes) were transiently transfected with the chosen combinations of cDNA constructs using the Lipofectamine/Plus method (Life Technologies Inc. (Gaithersburg, MD, USA)), following manufacturer’s instructions. Stably-transfected HeLa cells overexpressing either empty vector, GRK2-wt, GRK2_436-689_, or the mutant constructs GRK2-S670A or GRK2-K220R were described previously (Penela et al. 2010). HeLa-SX(p53) cells were transfected with empty vector, GRK2-wt or GRK2-K220R and antibiotic-selected cells were collected as pooled positive transfectants as reported (88).

### GRK2 knock-down

GRK2 knock-down was achieved by RNA interference using and adenovirus vector bearing the target GRK2 shRNA sequence (5’-GCAAGAAAGCCAAGAACAAGC-3’) using an un-related vector as control as previously described (88). Recombinant adenoviruses expressing the GRK2 shRNA were produced using BLOCK-iT™ Adenoviral RNAi Expression System (Invitrogen) according to the manufacturer’s instruction.

### cDNA constructs

The cDNAs encoding GRK2-wt, the kinase-dead mutant GRK2-K220R, the C-terminal construct GRK2_436-689_, the phosphorylation mutants GRK2-S670A, GRK2-Y86,92D (GRK2-Y2D) and GRK2-Y13,86,92F (GRK2-Y3F) have been previously described (22, 39, 88). The cDNAs encoding PI3K-p85 and PI3K-p110α were gently supplied by Dr. J.S. Gutkind (NIH, Bethesda, MD, USA). The construct coding the PIK domain (aa535-732) of PI3K-p110γ was provided by Dr. H.A. Rockman (Duke University Medical Center, Durham, North Carolina, USA). The ABL1 construct was from Dr. A.R. Nebreda (Institute for Research in Biomedicine, Barcelona, Spain). The constructs GST-GRK2_1-185_, GST-GRK2_185-467_ and GST-GRK2_436-689_ were previously described (Penela et al, 2010). The GRK2_1-147_ construct was kindly provided by Dr. J.L. Benovic (Thomas Jefferson Cancer Institute, Philadelphia, USA). The kinase-deletion mutants, GRK2-ΔNt (GRK2_Δ50-145_) were generously provided by Dr. C. Murga and GRK2-ΔCt (GRK2_Δ561-670_) was generated by PCR mutagenesis as described (36). The GRK2 construct mutated at the proline-box (GRK2-P2A; GRK2-P469,472A-R473A) was engineered by site-directed mutagenesis (in our laboratory and subcloned into the pcDNA3 vector) using the QuickChange® Lightning site-directed mutagenesis kit (Thermo Fisher) and the primers (forward 5’-CAAAAGTACCCTCCCGCGTTGATCGCCCCAGCAGGGGAGGTGAATGC-3’ and Reverse 5’-GCATTCACCTCCCCTGCTGGGGCGATCAACGCGGGAGGGTACTTTTG-3’) according to the manufacturer’s instructions.

### Reagents and antibodies

Recombinant PI3K-p85 regulatory subunit was obtained from Jena Bioscience GmbH (Jena, Germany). Affinity-purified rabbit polyclonal antibodies raised against GRK2 (C-15), the affinity-purified goat polyclonal antibody raised against actin (I-19), the anti-p21 goat polyclonal antibody (C-19), the affinity-purified anti-GST (B-14), anti-HA (F-7), and anti-p53 (DO-1) mouse monoclonal antibodies, the antiphosphotyrosine monoclonal antibody conjugated to horseradish peroxidase (PY99-HRP), the anti-MEK1 polyclonal antibody (C-18) and the polyclonal C-16 and C-14 antibodies that recognize ERK1 and ERK2, anti-α-tubulin (YOL 1/34) and anti-c-ABL (sc-56887) were purchased from Santa Cruz Biotechnology Inc. (Santa Cruz, CA, USA). The anti GAPDH (ab8245) was from Abcam. The was. The anti-GRK2/3 mouse monoclonal and the anti-PI3K-p85 rabbit polyclonal antibodies were purchased from Upstate Biotechnology (Lake Placid, NY, USA). The anti-pSer217/221-MEK1, anti-pTyr15-CDK1, anti-CDK1, anti-FKHRL1, anti-pThr32-FKHRL1, anti-pSer473-AKT, anti-AKT, and the anti-phospho-Thr202/Tyr204-ERK1/2 polyclonal antibody were purchased from Cell Signaling Technologies (Beverly, MA, USA). The anti-cyclin B1 monoclonal antibody was from BD Biosciences Pharmingen. Thymidine, aphidicolin, doxorubicin, and GW8510 were obtained from Sigma (St. Louis, MO, USA). LY294002 were supplied by Calbiochem (San Diego, CA, USA). Adaphostin was provided by Biaffin GmbH & Co KG (Kassel, Germany). Glutathione-Sepharose 4B beads were obtained from Pharmacia Amersham Biotech and G-protein sepharose from Invitrogen (Carlsbad, CA, USA. All other reagents were of the highest commercially-available grades.

### Cell cycle analysis and cell synchronization

HeLa cells and HeLa-SX(p53) were synchronized at G1/S transition using a thymidine-aphidicolin double block as described (22). Expression profiles of cyclin B, p-Y15-CDK1 and pS10-H3 were determined in cellular extracts by immunoblotting to confirm cell cycle progression of synchronized cells.

### Flow cytometry

The DNA content of cells growing asynchronously (1-1.5×10^6^) was determined by flow cytometry analysis. Cells were rinsed in cell cycle-staining buffer (0.1% sodium citrate, 0.3% Nonidet P-40, 50-μg/mL propidium iodide and 20 ng/mL RNase in PBS) and incubated for 30 min at room temperature. The fluorescent cells were detected with FACS Calibur (Becton-Dickinson) and data analyzed using the FlowJo Software (BD Biosciences).

### Immunoprecipitation and immunoblotting

For immunoprecipitation of GRK2 and co-immunoprecipitation of GRK2 with PI3K-p85 or ABL1 proteins, cells were lysed in 500 µl per 100-mm dish of lysis buffer A (Tris-HCl 20mM pH 7,5, 150mM NaCl, 1% Triton-X100, 0,1% SDS, 0,5% sodium deoxycholate) or B (50mM HEPES, pH 7.5, 150mM NaCl, 1 % Brij-96 V (FLUKA), 2mM EDTA, 1mM NaF, 1mM sodium orthovanadate), respectively. Upon centrifugation (15,000xg, 10 min), supernatants were incubated with a specific anti-GRK2 monoclonal antibody (clone C5/1.1, Upstate). Immune complexes were resolved in 7.5% SDS-PAGE and transferred to nitrocellulose membranes. After incubation with the indicated antibodies, blots were stripped and re-probed with a polyclonal antibody directed against the immunoprecipitated GRK2 protein (C15, Santa Cruz Biotechnology Inc). Blots were developed using a chemiluminescent method (ECL, Amersham). Band density was quantitated by laser densitometric analysis, and the amount of co-precipitated protein normalized to the amount of the immunoprecipitated protein, as assessed by the specific antibodies.

### GST pull-down assays

For the in vitro interaction of GRK2 with the PI3K-p85 subunit, GRK2-GST fusion proteins (100ng) were mixed with purified PI3K-p85 (500ng) in 50μl of binding buffer C (25mM HEPES pH 7.5, 100 mM NaCl, 1mM EDTA, 1% Brij-96V and 5% glycerol) and incubated for 90 min at 4°C before addition of glutathione-Sepharose 4B. Precipitated complexes were washed with the corresponding binding buffers and eluted with SDS sample buffer. Bound GRK2 or p85 was analyzed by immunoblotting using specific antibodies.

### Overlay protein interaction assays

For the SH3 domain-assay 15μg/μl of purified recombinant GRK2 protein generated in our laboratory, as previously described (22), were incubated with the SH3 domain-array II (Panomics, Cat #MA3011) following the manufacturer’s instructions. Hybridizations were visualized with anti-GRK2 (C-15, Santa Cruz) antibody and peroxidase-conjugated anti-rabbit secondary antibodies, followed by ECL plus (Amersham). Spots with intensities similar or stronger than positive controls (pos) were selected as SH3 domain-based ligands of GRK2.

### Clonogenic survival assays

Asynchronously growing cells (1-1.5×10^6^/100 mm-dish) were treated for 24h with different concentrations of doxorubicin and then plated at low densities in 6-well plates (200 cells/well) in triplicate. After 10 days of culturing in normal culture medium, the cells were fixed in methanol/acetic acid (3:1) solution and stained with Crystal violet for manual counting of live colonies. The surviving fraction (SF) was calculated with a correction for the plating efficiency (PE) as SF= PE (+doxorubicin)/ PE (-doxorubicin), and PE was determined as the ratio of the number of cell colonies to the number of plated cells.

### Statistical analysis

Data analysis was performed using GraphPad Prism for Windows. Means between groups were usually compared with unpaired Student’s t-test unless otherwise indicated, or by one-way ANOVA with Bonferroni’s or Tukeýs post hoc test, when indicated in the Figure legends. All results are expressed as mean±SEM.

