## SUPPLEMENTAL FIGURE LEGENDS for "GRK2-mediated AKT activation controls cell cycle progression and G2-checkpoint in a p53-dependent manner"

**^1^** Departamento de Biología Molecular, IUBM-UAM and Centro de Biología Molecular “Severo Ochoa” (UAM-CSIC), 28049 Madrid, Spain

**^2^** Instituto de Investigación Sanitaria La Princesa, 28006 Madrid, Spain

**^3^** Department of Molecular Oncology, Spanish National Cancer Research Centre (CNIO), 28029 Madrid, Spain

**^4^** CIBER de Enfermedades Cardiovasculares, ISCIII (CIBERCV), Madrid, Spain**^5^** **^5^**Lead contact

**Running title**: **GRK2-AKT and p53 networks in cell cycle regulation**

**SUPPLEMENTAL FIGURE LEGENDS**

**Fig. Supplementary S1. Expression of cell-cycle degradation-defective GRK2 mutants delays G2 progression into mitosis in cycling cells without altering the activation of the MAPK pathway**. In cells progressing synchronously in G2, the activation of ERK1/2 (A) and MEK1 (B) was unaffected by the presence of degradation-defective GRK2 mutants. Control HeLa, HeLa-A1, and HeLa-K1 cell cells were arrested in G1/S and released synchronously into the cell cycle, as detailed in Methods. The ERK1/2 and MEK1 activation was determined in whole-cellular lysates at the indicated times using an anti-phospho-Thr202/Tyr204-ERK1/2 antibody (p-ERK) and an anti-phospho-Ser217/221-MEK1 antibody (p-MEK1). The immunoblot was subsequently stripped, and total cellular ERK1/2 and MEK1 were detected with specific antibodies. A lower-mass form of phosphorylated MEK1 (p-MEK1∆) was observed. A similar-size band of MEK1 has been reported to correspond to an N-terminal proteolytic product of full-length MEK1, which is defective in ERK activation (Harding et al., 2003; Roberts et al., 2002). This band is identified by an anti-MEK1 antibody directed against the C-terminal region. Activation of MEK1∆ in each cell line tested increased at the G2/M transition (10h), peaking at mitosis (12h-14h), while the activation of ERK1/2 was less pronounced. (C) Both cyclin B upregulation and CDK1 activation were hindered at the G2/M transition in cells overexpressing GRK2-S670A or GRK2-K220R protein. Whole-cell lysates were analyzed by immunoblotting with anti-cyclin B and anti-pY15-CDK1 antibodies. After stripping, blots were probed with an anti-actin antibody as a loading control. Activation of CDK1 was detected as a decrease in inhibitory Y15 phosphorylation. In all panels, gels are representative of at least 3 independent experiments. Representative blots are shown.

**Fig. Supplementary S2. No increased activation of AKT occurs in the absence of GRK2 expression or without protein stabilization during G2 progression**. Cell lines stably expressing GRK2-wt (HeLa-WT5) (A), the competitor construct GRK2_436-689_ (B) which stabilizes the endogenous GRK2 protein (Penela et al., 2010), or an empty vector (HeLa-Ctrl), as well as HeLa cells transduced with an interference sh-GRK2 adenovirus construct or control adenovirus (C), were synchronized as detailed in Methods. Cell lysates at the indicated times were analyzed by immunoblotting with anti-pS473-AKT (p-AKT) and total AKT antibodies. Normalized p-AKT levels by total AKT were represented as fold-change with respect to maximal AKT activation in control parental (empty vector) cells (B). Data are the mean ± SEM of 3-4 independent experiments. The effectiveness of GRK2 RNA silencing was confirmed by immunobloting with anti-GRK2 antibody. Total Actin level was used as a loading control. Blots are representative of at least 3 independent experiments.

**Fig. Supplementary S3. GRK2 interacts with the SH3 domain of several SRC family tyrosine kinases, but its tyrosine phosphorylation in G2 is independent of c-Src or EFGR activities**. (A) Neither tyrosine phosphorylation of accumulated GRK2-S670A protein nor activation of AKT were affected by the inhibition of c-Src or EGFR kinase during the G2 phase. HeLa cells stably expressing GRK2-S670A (HeLa-A1) were arrested at G1/S and synchronously released into cell cycle in the absence or presence of c-Src inhibitor PP2 (5 µM) and EGFR inhibitor AG1748 (0.5 µM) added 2h post-release and maintained for the indicated time points. Tyrosine phosphorylation of immunoprecipitated GRK2 was analyzed as detailed in Methods, and cell lysates were analyzed by immunoblotting with anti-pS473-AKT (p-AKT) and total AKT antibodies. Representative blots of two independent experiments are shown. (B) Binding of GRK2 protein to the SH3 domain of c-Abl kinase in vitro. The Transignal SH3 domain-array (Panomics), consisting of 38 different SH3 domains immobilized on a membrane, was incubated with recombinant GRK2 protein (15 μg/μl), and GRK2-SH3 domain interacting pairs were immunodetected with a specific GRK2 antibody. Spots with an intensity 1.7-fold higher than the average internal positive signal were considered as reliable, and those with stronger intensities indicate higher binding affinity of SH3 domains to GRK2. A representative hybridized array membrane is shown with relevant targets coloured in red. ABL2, Abelson-Related Gene Protein; CRKL, CRK-like protein; EFS, Embryonal Fyn-associated substrate; NCK1, SH2/SH3 Adaptor Protein NCK-Alpha; OSF, Osteoclast stimulating factor I; PI3α, p85alpha subunit of PI3K; PIG2, 1-phosphatidylinos-4,5biphosphate phosphodiesterase gamma 2; RHG4, Rho-GTPase-activating protein 4; SNX9, Sorting nexin 9; c-Src, Tyrosine-Protein Kinase SRC-1; Tec, Tyrosine-protein kinase Tec.

**Fig. Supplementary S4. The absence of p53 neither affects the default cell cycle downmodulation of GRK2 in G2 nor its accumulation under genotoxic-induced G2 arrest**. (A) Knockdown of p53 expression in HeLa-SX(p53) cells in normal cycling and genotoxic-arrested conditions. Parental HeLa and HeLa-SX(p53) cells synchronously cycling after G1/S arrest were treated two hours after release with either vehicle or 1 µM doxorubicin for the indicated periods. Cellular lysates were subjected to immunoblot analysis with a specific p53 antibody. Actin levels were used as loading control. (B, C) Characterization of stable wild-type and mutant GRK2-overexpressing HeLa-SX(p53) cells. (B) Expression of GRK2 protein levels was analyzed in total lysates from p53-silenced cells stably transfected with mutant GRK2-K220R (HeLa-SX(p53)-K3s) and wt-GRK2 (HeLa-SX(p53)-WT7s) and compared to parental HeLa-SX(p53) cells and their counterparts in normal HeLa cells. (D) Effect of the absence of p53 on cell cycle regulated GRK2 protein levels. Synchronized cells as above with or without extra wild-type GRK2 (HeLa-SX(p53)-WT7s) or mutant GRK2-K220R (HeLa-SX(p53)-K3s) were exposed to doxorubicin or vehicle, and levels of total GRK2 and Actin were immunodetected with specific antibodies. Representative gels from two to three independent experiments are shown in all panels.

**Fig. Supplementary S5. Influence of p53 on GRK2-stimulated AKT during G2 checkpoint activation.** (A-C) Doxorubicin-induced stabilization of GRK2 lead to a less strong activation of AKT in the absence of p53 protein expression. (A) The p-AKT activation associated with the G2 checkpoint was lower in p53-deficient HeLa cells (HeLa-SX(p53)) compared to normal HeLa cells. Cells were exposed two hours post-release from a G1/S block to doxorubicin (1 μM) for the indicated times, and p-AKT and total AKT levels were analyzed by immunoblotting. (B-D) The activation of AKT in response to doxorubicin showed a tendency to increase with the upregulation of GRK2 in HeLa-SX(p53) cells. Representative blots (B) and quantification data (C) were shown. Synchronized p53-deficient cells with or without extra wild-type GRK2 (HeLa-SX(p53)-WT7s) or mutant GRK2-K220R (HeLa-SX(p53)-K3s) were exposed to doxorubicin or vehicle as above, and p-AKT and total AKT protein immunodetected. Levels of p-AKT normalized by total AKT in both cycling and doxorubicin-arrested cells were represented as fold-change with respect to 0h release (C). Data are mean ±SEM of 2-4 independent experiments. *** p< 0. 001 one-way Anova test when compared to 0 h post-release, and † p< 0.05, †† p< 0.01, ††† p< 0.01, one-way Anova test comparing with and without doxorubicin at each time point. When doxorubicin-exposed parental and stable GRK2 overexpressing cells were directly compared, the activation of AKT was clearly enhanced in cells expressing the degradation-defective GRK2-K220R mutant (D). Normalized pAKT data were represented as fold-change with respect to 4h doxorubicin-treated parental HeLa-SX(p53) cells (D). Data are mean ±SEM of 2-4 independent experiments. * p< 0. 05 one-way Anova test comparing with and without extra GRK2 at each time point, and ‡ p< 0.05, one-way Anova test when compared to 4 h Doxorubicin-treated HeLa-SX(p53)-WT7s cells. Representative blots are shown

**Fig. Supplementary S6. Doxorubicin-triggered GRK2 stabilization correlates with the induction of the G2-checkpoint response in the absence of p53.** Synchronized HeLa-SX(p53) cells, with or without extra wild-type GRK2 (HeLa-SX(p53)-WT7s) or mutant GRK2-K220R (HeLa-SX(p53)-K3s), were exposed to doxorubicin (1 µM) or vehicle two hours post-release from a G1/S block for the indicated times. Levels of Cyclin B1, phosphorylated H3 (pS10-H3), tubulin, and total and pY15-CDK1 proteins were immunodetected with specific antibodies. Cyclin B1 and pS10-H3 densitometry values were corrected by tubulin expression, and pY15-CDK1 levels were normalized by CDK1, represented as fold-change with respect to 0h release. Data are mean ± SEM of 2-6 independent experiments. * p< 0.05, when compared to doxorubicin-treated parental cells with HeLa-SX(p53)-WT7s and HeLa-SX(p53)-K3s cells at the corresponding times. † p< 0.05, †† p< 0.01, when compared to doxorubicin-treated with untreated HeLa-SX(p53) cells. § p< 0.05, §§ p< 0.01, §§§ p< 0.001, when compared to doxorubicin-treated with vehicle-treated HeLa-SX(p53)-WT7s cells. ‡ p< 0.05, ‡‡ p< 0.01, ‡‡‡ p< 0.001, when compared to doxorubicin-treated with vehicle-treated HeLa-SX(p53)-K3s cells. Representative blots are included.
